# Gempipe: a tool for drafting, curating and analyzing pan and multi-strain genome-scale metabolic models

**DOI:** 10.1101/2025.07.03.662949

**Authors:** Gioele Lazzari, Giovanna E. Felis, Elisa Salvetti, Matteo Calgaro, Francesca Di Cesare, Bas Teusink, Nicola Vitulo

**Affiliations:** Department of Biotechnology, University of Verona, 37134 Verona, Italy; VUCC-DBT, Verona University Culture Collection, Department of Biotechnology, University of Verona, 37134 Verona, Italy; Systems Biology Lab, A-LIFE, Institute of Molecular and Life Sciences (AIMMS), VU Amsterdam, 1081HZ Amsterdam, the Netherlands

**Keywords:** genome-scale metabolic models, strain-level metabolic biodiversity, bioprospecting

## Abstract

Genome-scale metabolic models (GSMMs) can mechanistically explain phenotypic differences among closely-related bacterial strains. However, high-throughput multi-strain reconstructions of GSMMs are still challenging: reference-based methods inherit curated information while missing new contents; alternatively, (universe-based) reference-free methods could cover strain-specific reactions, but they disregard curated information. Ideally, references should be curated pan-GSMMs for species (or genus), but their reconstruction is extremely demanding, making them still rare in literature.

Here Gempipe is presented, a computational tool streamlining the multi-strain reconstruction and analysis of GSMMs, going through the production of a pan-GSMM. Its reconstruction method is hybrid, as an optional reference GSMM is automatically expanded with extra reactions taken from a reference-free reconstruction. Gempipe also downloads, filters and annotates genomes; performs in-depth gene recovery; annotates models’ contents; predicts strain-specific capabilities. The companion programming interface includes functions ranging from the (pan-)GSMMs’ curation to the multi-strain analysis.

Gempipe was validated using multi-strain datasets, showing improved accuracy when compared with state-of-the-art tools. Moreover, metabolic diversities within *Limosilactobacillus reuteri* were explored, grouping strains into metabolically coherent clusters and systematically predicting health-related metabolites’ biosynthesis.

**IMPORTANCE:** Available GSMM reconstruction tools present major limitations in the context of multi-strain modeling. Gempipe surpasses these limitations by implementing a novel, hybrid reconstruction strategy. Not only it produces more accurate strain-specific GSMMs, but also pan-GSMMs when the only available reference is a manually curated model for a single strain, which is currently the most common case. With the vast availability of genome sequences, the high-throughput, multi-strain GSMM reconstruction and analysis approach provided by Gempipe will facilitate large-scale studies of the exploration and bioprospecting of strain-level bacterial metabolic diversity, moving a step forward in strains’ screening and rational selection.

## INTRODUCTION

Different strains of the same bacterial species can exhibit marked differences at the phenotypic level, such as the ability to catabolize different substrates, the presence of specific auxotrophies, or the acquisition of biosynthetic pathways through lateral gene transfer [1,2].

Genome-scale metabolic models (GSMMs) are systems-biology tools that describe the metabolic potential encoded by a genome. Assuming a steady state and specifying a biomass composition, their constraint-based simulations enable predictions of cellular growth under specific nutritive inputs [3]. Therefore, the availability of a GSMM for each strain of interest enables *in silico* screenings of phenotypic characteristics, offering a faster and more cost-effective alternative to traditional experimental methods, generating hypotheses to be subsequently validated experimentally.

Unfortunately, the creation of high-quality GSMMs is time-expensive because manual curation is required [4]. This bottleneck remains a key driver for the development of automated tools [5–9], some of which quickly gained traction due to their ability to produce simulation-ready GSMMs, using a reference-free, universe-based top-down approach [10].

Since the first pioneering work by Monk and colleagues on *E. coli* [11], the general steps for the multi-strain reconstruction of GSMMs remained mostly the same [12–18]. Briefly, genomes are collected and filtered for quality; then, genes are predicted and clustered, creating orthologous gene families sometimes referred to as the pangenome; one of the strains includes a high-quality, manually curated GSMM that is used as reference; for each strain, a copy of the reference is made and all the genes that do not have an ortholog with the reference are subtracted, consequently removing associated metabolic reactions; finally, gap-filling is usually limited to minimal media as the starting genomes are quality-filtered and excessive gap-fillings could hide true strain-specificities.

The above reference-based method was formalized in 2019 by Norsigian and colleagues [19], where metabolic functions are inherited from the reference GSMM after orthologous genes are detected via a blastp [20] best reciprocal hits (BRH) alignment. The method was then implemented in Bactabolize [21], a recent tool published in 2023. Overall, its efficacy is dependent on the availability of a curated and phylogenetically close GSMM taken as reference.

However, the method has a key limitation: output GSMMs contain subsets of the reactions in the reference, excluding unmodeled strain-specific reactions. Indeed, the protocol [19] requires a manual curation of the output GSMMs, adding new reactions that were not originally in the reference, a step that was not automatized in Bactabolize [21]. For this reason, to fully capture strain-specific metabolic features, a curated pan-GSMM that encompasses the metabolic diversity of the entire species (or genus) should be provided as reference instead of a strain-specific GSMM.

However, while curated strain-specific GSMMs are time-consuming to produce and therefore often lacking for non-model organisms, comprehensive pan-GSMMs are even more challenging to obtain. When manually curated, pan-GSMMs can require years of development [16,22,23] and indeed they are still rare in literature [21]: the few covered species include *Klebsiella pneumoniae* [23], *Escherichia coli* [24], *Salmonella enterica* [12] and *Bacillus subtilis* [17,18].

Given the vast number of strain-specific genome sequences now available and the scarcity of comprehensive pan-GSMMs, there is a growing need for tools performing multi-strain reconstructions efficiently, even in the absence of a pan-GSMM. These tools should not only be capable of using a reference strain as a starting point, but also of autonomously integrating new, strain-specific reactions to fully capture the metabolic diversity across strains.

In this work Gempipe is introduced, a novel package that, to the best of our knowledge, is the first to offer pan and multi-strain reconstruction of GSMMs by implementing a hybrid reconstruction method where an optional reference GSMM is automatically expanded with new contents transferred from an independent reference-free reconstruction. Gempipe also provides additional features: retrieval and quality-filtering of genomes; gene annotation; in-depth gene-recovery; re-annotation of modeled contents; a companion application programming interface (API) helping the manual curation of (pan-)GSMMs; an “autopilot” mode skipping the (recommended) manual curation; various flux-balance analysis (FBA)-based predictions of strain-specific metabolic features; dedicated API functions for the multi-strain analysis. In this sense, Gempipe is not only a reconstruction tool, but also an analysis tool in the context of biodiversity exploration/bioprospecting.

## MATERIAL AND METHODS

### From genomes to gene clusters

Gempipe is composed of three command-line programs, “gempipe recon”, “gempipe derive”, and “gempipe autopilot”, along with a Python API (**Figure 1**). “gempipe recon” reconstructs draft pan-GSMMs and supports four types of inputs: proteomes in genbank format, proteomes in FASTA format, genome assemblies in FASTA format, and a list of NCBI Species Taxonomy IDs (taxids). When taxids are given in input, all the available assemblies for the indicated species are automatically downloaded from Genbank. Each genome/proteome is treated as a separate strain.

**Figure 1.**
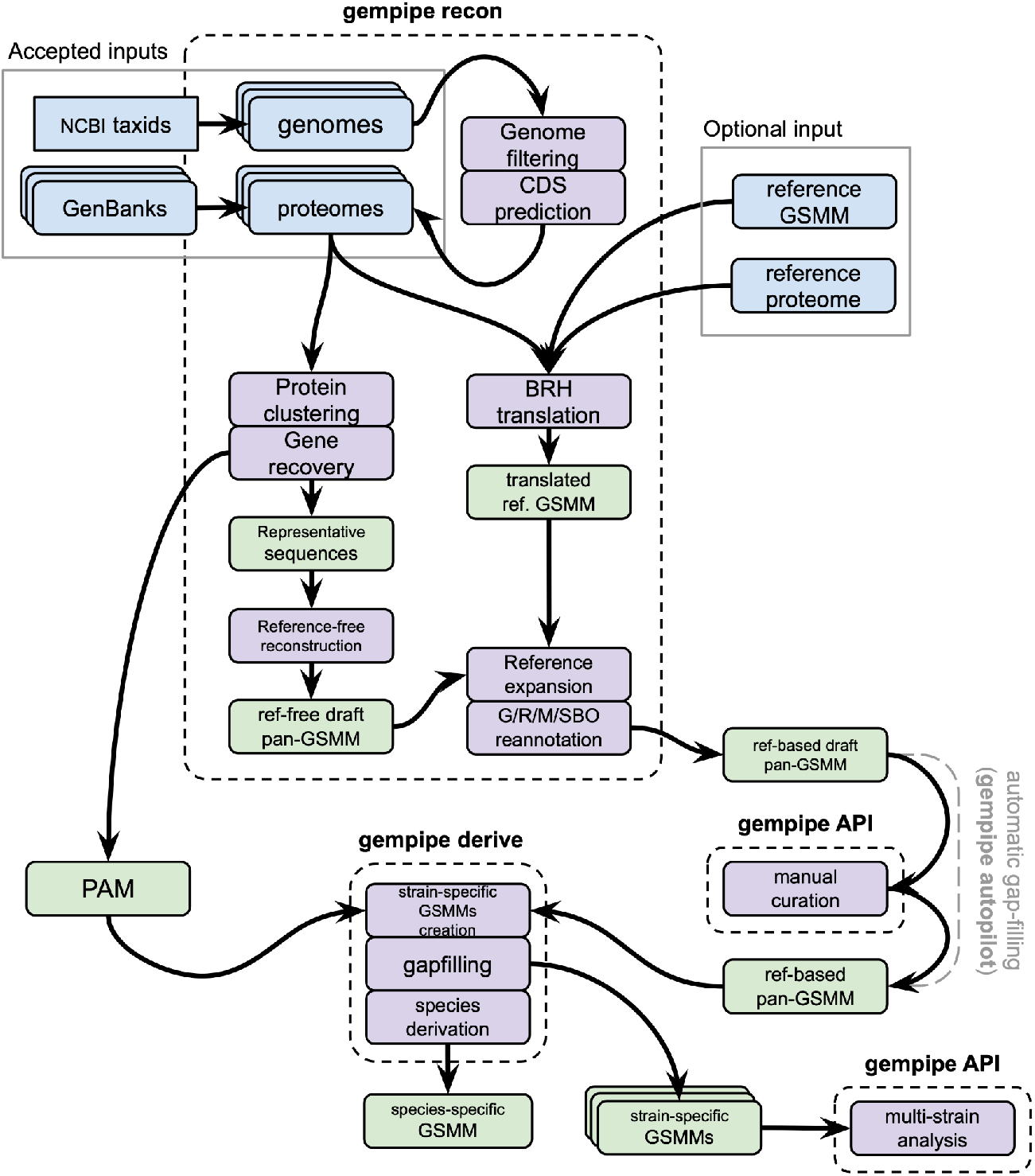
Overview of Gempipe. “gempipe recon” creates a draft pan-GSMM and the presence-absence matrix (PAM), containing the information relative to the protein clustering; “gempipe derive” derives strain-specific GSMMs, starting from the PAM and the pan-GSMM; the latter should be manually curated beforehand, e.g. by using dedicated functions of the Gempipe API. When using “gempipe autopilot”, the manual curation is substituted by an automatic gap-filling of the draft pan-GSMM. Finally, the Gempipe API can be used once again to perform multi-strain analyses. When strains of different species are inputted in the same run, species-specific GSMMs are also produced, defined by the set of reactions always present in all strain-specific GSMMs belonging to the same species.

When genomes are inputted, they are first subjected to gene annotation running Prodigal v2.6.3+ [25] via Prokka v1.14.6+ [26] to obtain conveniently named proteomes. Next, BUSCO v5.4.0+ [27] is run indicating the database of interest, which is automatically downloaded. Subsequently, *seqkit stats* v2.2.0+ [28] is run to compute the number of contigs and N50 for each assembly. After these steps, strains that do not fulfill all the following user-specified thresholds are discarded from subsequent analysis: maximum number of missing (default 2%) or fragmented (default 100%) BUSCO orthologs, minimum N50 (default 50000 [29]), and maximum number of contigs (default 200 [29]).

When a proteome is obtained for each strain, amino acid sequences are grouped into clusters based on high global sequence identity (90%). This is done by using CD-HIT v4.8.1+ [30] with parameters *-M -g 1 -aL 0*.*70 -aS 0*.*70 -d 0 -c 0*.*90*, obtaining a representative sequence for each cluster. Using the clustering information, an initial gene presence / absence matrix (PAM) is created, with the cluster IDs in rows, strains in columns, IDs of strain-specific member genes in cells.

Next, a three-step gene recovery is applied to mitigate possible errors arising from genome assembly or gene calling. Three scenarios are addressed (see **Supplementary Information 1.1.1**): (1) premature stop codon that breaks a protein sequence in two pieces; (2) sequences located in genomic regions overlooked by the gene caller; (3) sequences overlapping [31] a previously annotated gene and an overlooked region. The PAM is updated with the recovered sequences. When proteomes are inputted, they are directly used in subsequent analysis and no gene recovery is performed.

### Reference-free draft pan-GSMM generation

The representative sequences of clusters are processed to create a draft pan-GSMM using a reference-free reconstruction approach. This pan-GSMM is used to expand the optional reference GSMM with new strain-specific contents; alternatively, when no reference GSMM is provided, it is directly used in downstream analysis.

This reference-free reconstruction phase is based on the bacterial universes (Gram positive or negative) provided by CarveMe v1.5.2 [5]. Even if Gempipe and CarveMe share the same universes (and underlying gene database), the reference-free reconstruction algorithm differs: Gempipe is more conservative and accounts for different gene isoforms while preserving the original enzyme complexes as defined in BiGG [32], a database of manually curated GSMM, leading to enhanced gene-to-reaction association (GPR) rules (see **Supplementary Information 1.1.2**).

### Expanded reference-based draft pan-GSMM generation

When an optional reference GSMM is provided (together with its associated proteome), it is used as the cornerstone of the reconstruction process. First, reference gene IDs are translated into cluster IDs, making the reference GSMM compatible with the PAM and the previously made reference-free draft pan-GSMM. This translation is based on orthologs determination via blastp [20] BRH alignments between each strain and the reference proteome, similarly to the approach suggested by [19] (see **Supplementary Information 1.1.3**).

The translated reference GSMM is then expanded with new gene clusters, reactions and metabolites taken from the reference-free draft pan-GSMM used as a repository of new contents, leading to the production of an expanded reference-based draft pan-GSMM. The latter inherits from the reference key features such as the non-growth associated maintenance energy (NGAM) [4] and the biomass equation [33], which would be otherwise inherited from the CarveMe universe [5]. Moreover, during the expansion phase, curated information contained in the reference GSMM in terms of metabolites’ mass/charge and reactions’ balancing, is respected (see **Supplementary Information 1.1.4**).

The final draft pan-GSMM is subjected to reannotation of its metabolites, reactions, genes, and Systems Biology Ontology (SBO) terms [34]. This facilitates prospective uses of the output GSMMs, and can lead to better scores in the community standard test-suite MEMOTE [35]. This reannotation is mainly (but not exclusively) based on MetaNetX v4.4 [36] (see **Supplementary Information 1.1.5**).

### Derivation of strain-specific GSMMs

Once the pan-GSMM has been sufficiently curated (the Gempipe API can be used for this task, see **Supplementary Information 1.1.6**), it is inputted into “gempipe derive” together with the PAM, producing a strain-specific GSMM for each strain. Briefly, for each strain, a copy of the pan-GSMM is made. If the strain has no genes in a cluster, the cluster is removed, potentially leading to the loss of associated reactions. Similarly, if all genes in a cluster have premature stop codons, the cluster is removed. Next, reactions are iterated while updating their GPR: each remaining cluster ID is replaced by the corresponding strain-specific genes.

Each strain-specific GSMM is then gap-filled using a user-provided recipe for a medium, preferably minimal, known or assumed to support growth of all the input strains. More than one recipe can be provided, leading to multiple rounds of gap-filling. If no media file is provided, a generic minimal aerobic medium recipe is used, having glucose, ammonia, phosphate and sulfate and as sole C, N, P and S sources, respectively. The COBRApy [37] gap-filling algorithm is applied, using the pan-GSMM as source of reactions and a user-selectable minimum flux through the objective reaction. Moreover, the strain-specific gap-filling step can optionally be skipped, which is useful for example when auxotrophies has to be studied on minimal media [24]. At this point, strain-specific GSMMs have the minimum requirements to be used in simulations.

Before proceeding with “gempipe derive”, the draft pan-model should be curated, e.g. by using dedicated functions of the Gempipe API. As an alternative, strain-specific GSMMs can be seamlessly produced from genomes/proteomes by using “gempipe autopilot”, which skips the manual curation by applying an automated gap-filling to the draft pan-GSMM. This gap-filling is prioritized by using penalties derived alignment metrics (see **Supplementary Information 1.1.7**).

### Multi-strain predictions and analyses

Once obtained the strain-specific GSMMs, in addition to the Biolog® PM simulations, specific metabolic features can be predicted, including: capability to catabolize alternative C, N, P or S substrates, presence of auxotrophies for amino acids and vitamins, and potential biosynthesis of specific metabolites (see **Supplementary Information 1.1.8**). These features, together with the presence of reactions in strains, are stored as binary feature tables (BFTs). These tables have strains in columns, binary features in rows, and 1 (feature presence) or 0 (absence) in cells.

The Gempipe API contains dedicated functions for multi-strain analysis, where any number of BFTs can be inputted. Briefly, BFTs are combined into a single table and the pairwise similarity between strains is then calculated using the Jaccard index. This produces a distance matrix, which is further processed to create a dendrogram using Ward’s agglomerative clustering [11]. The latter is referred to as a “phylometabolic tree”, where strains with similar metabolic potential are placed closely. Clusters of metabolically coherent strains can be extracted from a phylometabolic tree and their characteristic features identified (see **Supplementary Information 1.1.9**). Tutorials for multi-strain analyses using the Gempipe API are available in the Gempipe documentation.

## RESULTS

### Models’ contents and similarity between tools

To validate the multi-strain reconstruction of Gempipe, three strain-specific dataset were used, named “*Klebsiella*”, “*Ralstonia*” and “*Pseudomonas*”, composed by 37, 11, and 36 strains, respectively (see **Supplementary Information 1.2.1**). Comparisons were made with current state-of-the-art reference-free reconstruction tools, namely CarveMe [5] and gapseq [7], as well as a recent reference-based tool focused on strain-specificity studies, Bactabolize [21] (see **Supplementary Information 1.2.2**). Comparisons were focused on automation, therefore manual curation was skipped and “gempipe autopilot” was used.

Contents of reconstructed strain-specific GSMMs were compared against their relative manually curated reference (**Figure 2A**). Moreover, every tool was compared against each other using the mean Jaccard index of the reaction content (**Figure 2B**) (see **Supplementary Information 1.2.3**).

**Figure 2.**
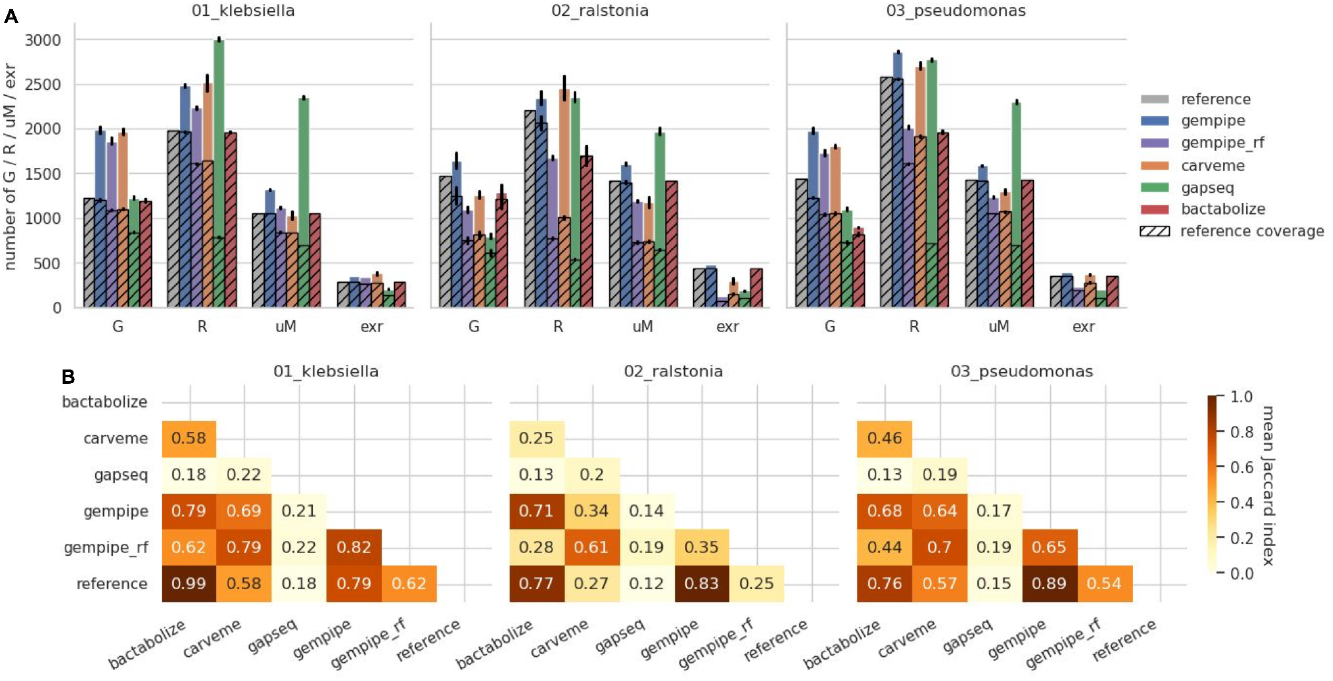
Comparison of the general reconstruction metrics. “gempipe_rf” indicates Gempipe ran without reference. **A)** Content comparison. ‘G’: number of genes; ‘R’: number of reactions excluding exchange reactions; ‘uM’: number of unique metabolites, i.e. not considering their compartment; ‘exr’: number of exchange reactions. Bar height corresponds to the mean between strains, while error bars represent standard deviations. Hatched area represents contents in common with the reference. **B)** Similarity between tools based on reaction content. Cells report the mean Jaccard index along the strains, computed for the reaction IDs.

Leveraging its hybrid reconstruction method (expanded reference-based), Gempipe had a generally better reference coverage than the other tools, including purely reference-based tools (Bactabolize). On the other hand, output models went beyond the reference with a higher number of modeled contents, aligning with the reference-free tools. The mean Jaccard index confirmed a better reference coverage for Gempipe, where the apparent lower performances in the *Klebsiella* dataset are only due to the addition of new reactions during the reference expansion phase. When run without a reference, Gempipe’s models were more similar to the ones produced by CarveMe, likely due to the shared BiGG-based database. However, the introduction of reactions and metabolites in Gempipe is clearly more conservative when using default parameters. gapseq models, despite having a consistently higher number of metabolites, were the most divergent from the reference. However, the conversion between SEED and BiGG IDs provided by MetaNetX [36] is not perfect, so the coverage metrics reported for gapseq could have been underestimated.

### Phenotype prediction accuracy

To evaluate the ability to recapitulate phenotypic traits, publicly available binarized Biolog® PM data were used as a benchmark (**Figure 3, Supplementary Table 1**), where the kinetic signal was converted into a binary response “can grow” / “cannot grow”. The cases of no growth and infeasible solution were distinguished, and the latter penalized (see **Supplementary Information 1.2.4**).

**Figure 3.**
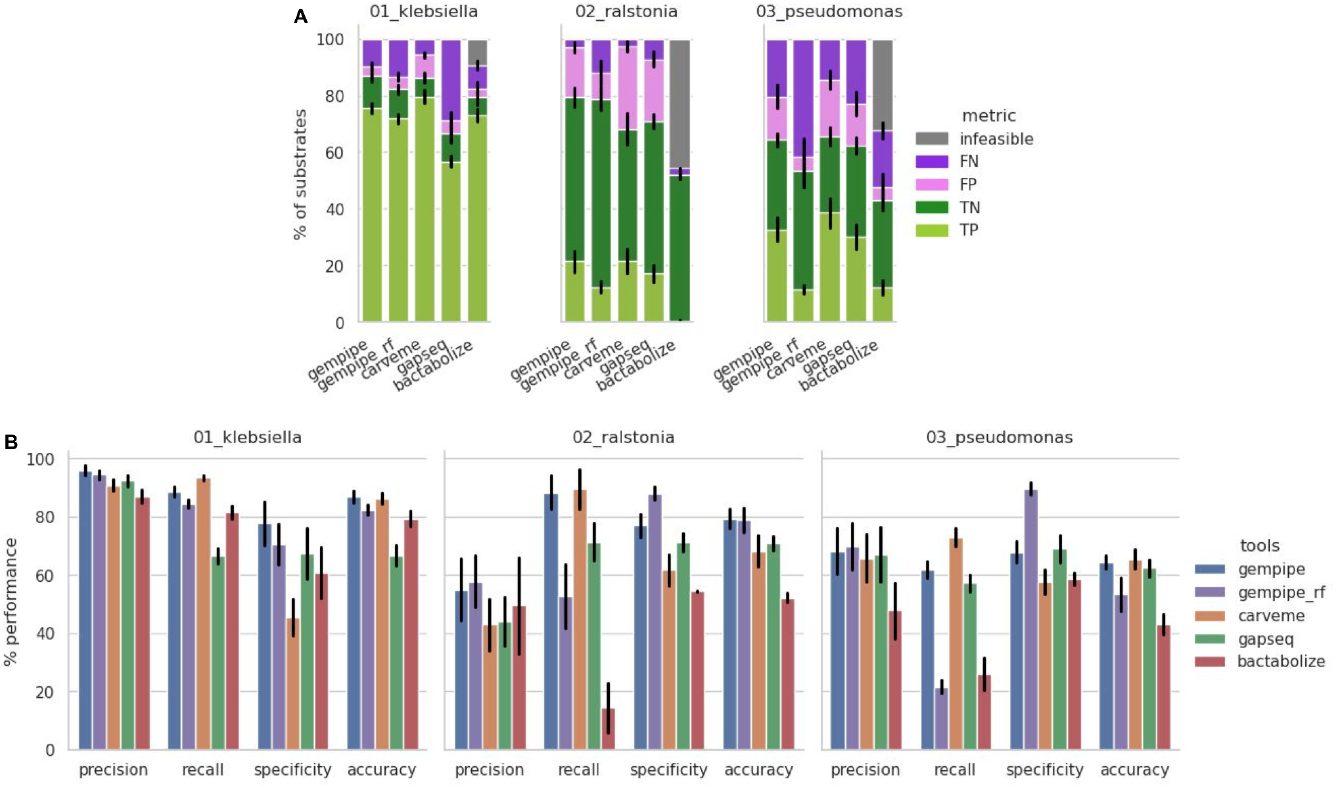
Comparison between experimental and simulated Biolog® PM growth assays. Bar height corresponds to the mean between strains, while error bars represent standard deviations. “gempipe_rf” indicates Gempipe ran without reference. **A)** Outcome of the comparison considering single substrates. TP: true positive; TN: true negative; FP: false positive; FN: false negative; infeasible: FBA without solution. **B)** Overall metrics for the substrate utilization prediction. Infeasible simulations have been penalized.

In general, the accuracy of Gempipe was better or in line compared to the other tools (**Figure 3B**), particularly when using its hybrid reconstruction mode. While CarveMe was close to Gempipe in terms of mean accuracy, it was observed that its internal gap-filling algorithm, designed to *“enforce network connectivity”* [5], tended to maximize the number of substrates for which growth is predicted; this led not only to fewer FNs (**Figure 3A**), resulting in better recall (**Figure 3B**), but also to more FPs, resulting in detrimental specificity. Despite using the same CarveMe’s assets (gene database and reaction universes), the different implementation of Gempipe better represented the no-growth phenotypes.

Benefits are even more evident when modeling genera still without reference GSMMs deposited in BiGG [32], as they cannot be represented in CarveMe’s assets, like e.g. *Ralstonia*. Bactabolize failed to recover orthologs for many reference genes, resulting in highly gapped reaction networks, despite the medium-specific gap-filling step in common with the other tools. As a consequence, many FBAs performed during Biolog® simulations resulted “infeasible”, preventing the usage of this purely reference-based tool, in particular for the *Ralstonia* and *Pseudomonas* datasets.

### Remaining orphan reactions

The quality of reconstructions may also be evaluated by the number of modeled metabolic reactions (not exchanges, sinks, nor demands) which have not been associated with genes and, at the same time, have not been labeled as spontaneous. These reactions, also known as “orphan” reactions [38], can be left by internal gapfillers of automated reconstruction tools to improve network connectivity [5]. Ideally, the number of orphans should be minimized by manually checking and reassociation to the corresponding genes: a high number of orphans may indicate insufficient curation and excessive reliance on gap-filling.

The presence of orphan reactions was compared (**Figure 4**), and Gempipe reconstructions contained the lowest fraction in every dataset. In Bactabolize reconstructions, orphans were copied from the reference, but the fraction in *Ralstonia* and *Pseudomonas* datasets was inflated due to the gap-filling. In Gempipe, orphans are also copied from the reference but, during the reference expansion phase, their GPRs are supplemented with missing gene clusters taken from the independent reference-free reconstruction. This resulted in a fraction of orphans lower than the reference in all the three dataset, and even lower with respect to the reference-free tools. Respect to CarveMe, in particular, the difference seemed to be more accentuated when the reference was not part of the BiGG collection [32], as in the *Ralstonia* dataset. When Gempipe was run without a reference, the number of orphans reached its minimum; in this reconstruction mode, remaining orphans are consequence of the two biomass-centered gap-filling steps: the first applied to the draft pan-GSMM, and the second to the strain-specific GSMMs.

**Figure 4.**
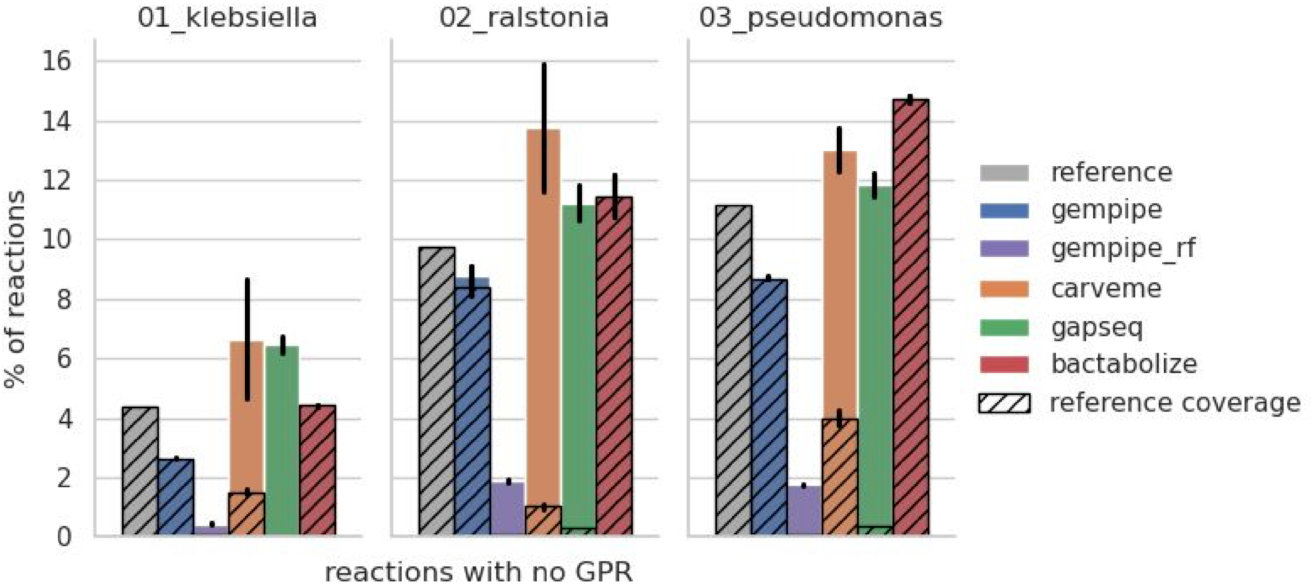
Relative number of modeled reactions with no GPR (orphans). Exchange, sink, and demand reactions are excluded, as well as reactions containing the substring “diffusion” in their name. Bar height corresponds to the mean between strains, while error bars represent standard deviations. Hatched area represents orphans in common with the reference.

### Metabolic biodiversity of *Limosilactobacillus reuteri*

*Limosilactobacillus reuteri* (*Lr*, formerly *Lactobacillus reuteri*) is a species of lactic acid bacteria (LAB) adapted to the gastrointestinal tract (GIT) of vertebrates and widely studied for its probiotic potential [39]. Recently, 6 subspecies of *Lr* were formally proposed by Li and colleagues [39], and characterized both phylogenetically and phenotypically. Subspecies reflect the host: *Lr* subsp. *reuteri* is adapted to human and herbivores; *Lr* subsp. *kinnaridis* to humans and poultry; *Lr* subsp. *porcinus* and *Lr* subsp. *suis* to pig; *Lr* subsp. *murium* and *Lr* subsp. *rodentium* to rodents [39].

Gempipe was used to explore the metabolic biodiversity of *Lr* at the strain level. 1056 *Lr* assemblies, of which 597 derived from metagenomes (MAGs), were retrieved. Contigs belonging to other species were removed from MAGs (see **Supplementary Information 1.3.1**). The 545 genomes remained after taxonomy and quality filtering (**Supplementary Table 2**) were assigned to subspecies based on ANI thresholds reported in [39]: 69 resulted classified as *kinnaridis*, 63 as *reuteri*, 52 as *rodentium*, 21 as *suis*, 2 as *porcinus*, 115 as *murium*, while the remaining were not classified (**Supplementary Figure S1**).

A curated GSMM for *Lr* JCM1112 [40] was used as reference in Gempipe (see **Supplementary Information 1.3.2**). Due to the hybrid reconstruction mode, it was expanded with new strain-specific contents generating a draft pan-GSMM. From the latter, 545 GSMMs were derived. BFTs for metabolic reactions, auxotrophies, and growth on alternative C sources were used to build a phylometabolic tree (**Supplementary Figure S2**). Since vitamin B12 is unrelated to the ecological niche/subspecies [41], the 24 reactions forming the B12 biosynthetic pathway were removed before generating the tree. A number of clusters equal to the subspecies (6) was extracted. Clusters were generally consistent with the subspecies: *reuteri* and *porcinus* were substantially contained in Cluster_3, *kinnaridis* in Cluster_5, *suis* in Cluster_1, *murium* and *rodentium* in Cluster_2 (**Figure 5A**).

**Figure 5.**
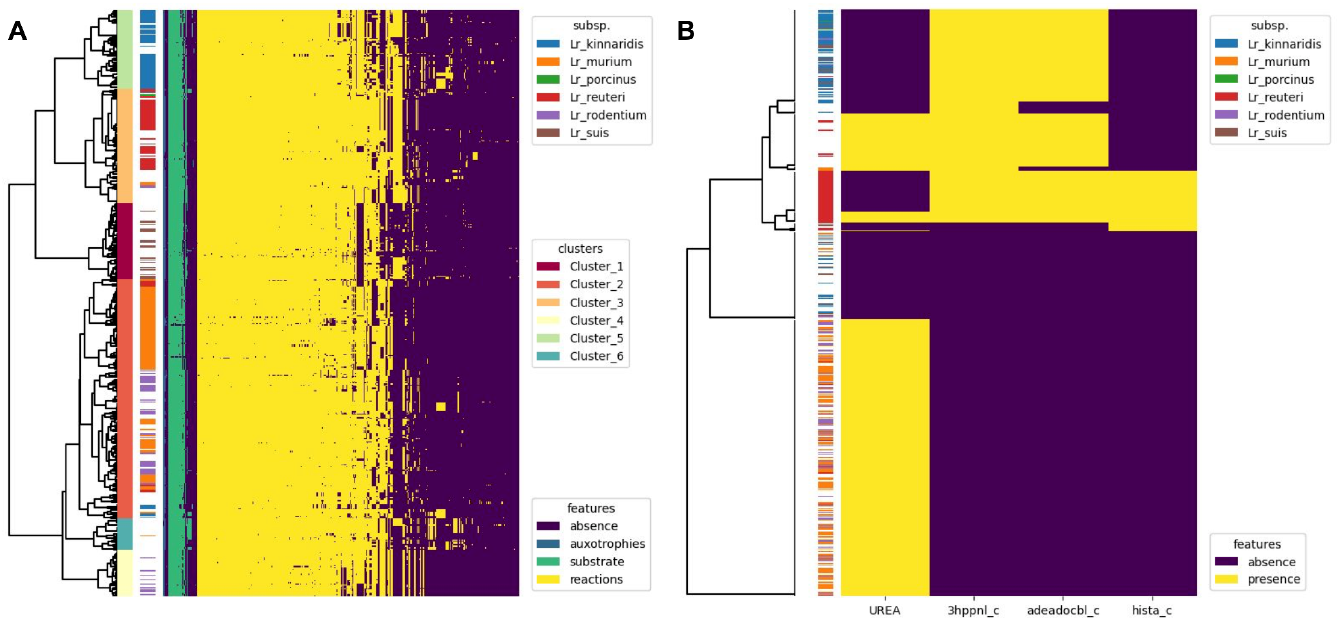
Multi-strain analysis using the Gempipe API functions. **A)** Phylometabolic tree built using presence/absence data for reactions, auxotrophies, and growth on alternative C substrates. 6 clusters of metabolically coherent strains were extracted from the tree and shown alongside the subspecies attribute. Only features not constant across strains are represented. **B)** Potential strain-specific production of health-related metabolites vitamin B12 (“adeadocbl_c”), reuterin (“3hppnl_c”), and histamine (“hista_c”), and presence of urease (“UREA”). The subspecies attribute is reported.

Consumption data for 49 C substrates characterizing the 6 subspecies were reported by Li *et al*. [39] (“general” dataset). These data were derived by the same authors from experiments on 12 strains, 2 for each subspecies (“specific” dataset) (**Supplementary Table 3**). The “specific” dataset was compared with strain-specific simulations, which resulted in 93% accuracy (mean computed on 11 strains, as GCA_000712565.2 was quality-filtered). The “general” dataset was compared with the feature’s relative frequency within clusters (see **Supplementary Information 1.3.3**); this comparison revealed interesting discrepancies between literature and modeled phenotypes.

For example, growth on galactose is reported as species capability [39]; however, ∼8% of strains (45), mainly belonging to *murium* and *rodentium* subspecies, were predicted unable to catabolize this substrate. Upon further investigation, these strains were consistently lacking the aldose 1-epimerase, possibly explaining the deficient phenotype (see **Supplementary Information 2.1**). Among compared substrates, D-xylose was particularly interesting for three reasons: (1) its catabolic route was not in the reference GSMM, but it was automatically included by Gempipe during the reference expansion phase; (2) its utilization was predicted to have high variability across strains, ranging from ∼9% in *murium* to ∼88% in *rodentium*; (3) its simulation accuracy with the “specific” dataset was 100%. For instance, only ∼38% of the 21 strains strictly classified as *suis* were predicted to grow on D-xylose, while this is reported to hold for the entire subspecies [39]. This further corroborated the idea that phenotypic descriptions of species/subspecies provided in literature are actually not always valid, possibly because they are based on a number of strains (2 in this case) too low for generalization, or due to limited reliability of phenotypic testing. In this context, the GSMM-based analysis gave more comprehensive results than traditional phenotypic characterisation.

In humans, *Lr* is found in many sites other than GIT, including breast milk, skin, and urinary tract. *Lr* is reported as probiotic, and some of the metabolites it produces have a proven health-related effect [42,43]. For example, reuterin (mixture of different forms of 3-hydroxypropionaldehyde) is released by several strains and exerts antimicrobial activity against Gram negative bacteria, making *Lr* effective against GIT infections [42,44]. Histamine is another strain-specific metabolite, acting as intestinal immunomodulator and anti-inflammatory agent [42,44]. Vitamin B12 (cobalamin) is an essential vitamin in humans, introduced with the diet; only four B12-producing *Lr* strains were clearly identified as of 2018 [44]. Apart from health-related metabolite production, other metabolic features are of interest to assess adaptation strategies in different hosts; one of them is the conversion of urea to ammonium and CO_2_ (urease), likely involved in acid resistance [45].

Gempipe enabled a systematic evaluation of these four key metabolic features in all the 545 filtered strains (**Figure 5B**). The majority of rodent-associated strains (*Lr* subsp. *murium* and *rodentium*) were predicted capable of urease activity, probably needed for survival in rodent GIT [46,47]; this is in accordance with a recent genome-wide association study (GWAS) [48]. Moreover, most but not all of the human- and poultry-associated strains were predicted as potential producers of reuterin and vitamin B12, with the underlying genes being indeed part of the same *pdu-cbi-cob-hem* gene cluster [46]. Interestingly, histamine production was an exclusive feature of subsp. *reuteri* [39], with only a few *reuteri* strains lacking this trait.

## DISCUSSION

In the present work Gempipe was introduced, a multi-purpose package for pan and multi-strain genome-scale metabolic modeling. It adopts an hybrid reconstruction approach that lies between reference-free and reference-based methods, as an optional reference is expanded with new contents coming from a universal GSMM. Together with an internal clustering of strain-specific genes, an effective in-depth gene recovery (see **Supplementary Information 2.2**), and conservative generation of GPRs, the implemented approach proved to be effective for multi-strain reconstructions, with better or similar performances compared to current established reconstruction tools, when focusing on metabolic features without considering manual curation.

Gempipe represents a third option in the panorama of reconstruction tools: reference-based methods use a manually curated model (or sorted list of models, from the phylogenetically closest to the most curated) to be used as template, from which reactions are copied after orthologs genes are determined [49–51]; instead, reference-free methods use a semi-curated universal model with a generic biomass equation, from which reactions are copied and gap-filled based on sequence homology (one-way alignment) [5,7]. From the first approach, Gempipe inherits the ortholog determination to translate reference genes into equivalent gene clusters; from the second, Gempipe inherits the homology-based insertion of new, strain-specific reactions, with GPRs already based on gene clusters.

In the case of multi-strain reconstruction, reference-based methods are usually applied [11–18]. The main concern, in this context, is that the template must be representative of the metabolic diversity of the entire species (or genera) [21], otherwise strain-specific reactions must be added afterwards to each generated model [19]. This is why a comprehensive (pan) GSMM is usually curated prior to running reference-based reconstruction tools [23]. However, such pan-GSMMs representing a species (or genus), are complex to build and still rare in literature [12,18,23,24], with as little as 4 species reported [21]. Indeed, in some GSMM-based biodiversity studies, strain-specific models have been used as reference [15–17]. This is a limiting approach, as generated models will just be a subset of a single strain. In this context, Gempipe was able to grasp strain-specificities better than purely reference based methods like Bactabolize [21] when a strain-specific GSMM is used as reference instead of a pan-GSMM, which is a common case. Moreover, inheriting contents from a manually curated reference, the models generated by Gempipe provided more accurate predictions with respect to those built with reference-free methods like CarveMe [5] and gapseq [7]. All this was achieved while minimizing the number of orphan reactions, and maximizing the MEMOTE metrics [35] (see **Supplementary Information 2.3**).

Considering their paucity, Gempipe emerges also as a valuable tool to quickly build and curate pan-GSMMs to be used in biodiversity or bioprospecting studies. It must be noted, however, that the concept of pan-GSMM was introduced also in the context of metagenomics-derived models [52], which is remarkably different. In this context, indeed, pan-GSMMs are instrumental to cope with the incompleteness and contamination of MAGs, and strain-specificity (in terms of genes and reactions) is lost in favor of a consensus/mean reconstruction representing a species-level genome bin (SGB) [52]. Instead, in the GSMM-guided exploration of biodiversity, the context where Gempipe operates, strain-specificity must be retained and emphasized as it is needed for the subsequent generation of strain-specific GSMMs [19].

In the development of Gempipe, particular attention was put into how metabolic features are encoded in the model. In this regard, it must be noted that metrics commonly used to evaluate and compare reconstructions (accuracy, precision, recall, specificity) [5,7,21] do not take into consideration the faithfulness of the reaction network in representing the organism. Indeed, behind a true positive match with the experimental data (e.g., Biolog® screenings) there could be cases of (i) wrong reaction mechanisms (e.g., erroneous transporter types); presence of metabolic reactions not supported by genes (orphans); (iii) reactions with seriously impaired GPRs, lacking many components of a protein complex. Provided that manual curation still remains essential for obtaining truthful GSMMs, users should be aware of the reconstruction principles that each tool follows to draft a model. Gempipe includes reactions only when a protein complex is fully supported by genes, otherwise mismatches (false negatives) will lead the manual curation in closing the gaps. On the other side, tools providing (a) internal gap-filling procedures aimed to improve the network connectivity and (b) too permissive mechanisms of GPR generation and reaction inclusion, could lead not only to predict a higher number of growth-supporting substrates (higher recall and lower specificity) but, most importantly, to match the experimental data with a biologically inaccurate representation of the metabolism, which can be easily overlooked during the manual curation.

Finally, Gempipe is not meant to be just a reconstruction tool, but also an analysis tool in the context of biodiversity exploration. Indeed, while the Gempipe API includes functions to curate models, it also contains functions to analyze the deck of strain-specific GSMMs in output. For example, functions are included to cluster strains according to their metabolism, and to visually compare metabolic clusters with respect to other attributes, such as niche metadata or the formal species or subspecies classification. This aids users to achieve goals including (i) the screening of strains for desired metabolic traits, (ii) the classification of strains according to their metabolic capabilities, and (iii) the definition of species and subspecies, and possibly other taxonomic ranks, in terms of their core metabolic potential. In this context, the case study on *L. reuteri* here reported provided insights also from a taxonomic point of view (see **Supplementary Information 3.1**).

Limitations of Gempipe are mostly due to the resources it relies on. The BiGG namespace [53] was adopted as it was convenient for two main reasons: (i) the availability of high-quality, manually curated reference models based on the same namespace [32]; (ii) the human readable IDs, particularly useful during manual curation, when GSMM-based metabolic maps have to be hand-drawn or interpreted [54,55] (for example, D-glucose is “glc D”, immediately recognizable with respect to its ModelSEED [56] equivalent “cpd00027”). While convenient, the BiGG database has imperfections due to its structure: it is not a coherent biochemical database, but rather a collection of models from which content a biochemical database is derived. Depending on the model of origin, the same metabolite or reaction can be defined differently. Consequences are many: (i) the same reaction can be represented with different reversibility (e.g. “<u>ILETA2</u>“); (ii) the same metabolite can be represented with different IDs, leading to duplicate metabolites (e.g. “<u>ind_c</u>“ / “<u>indole_c</u>“); (iii) the same reaction can use duplicate metabolites, leading to duplicate reactions (e.g. “<u>TRPS2</u>“ / “<u>TRPS2_1</u>“); (iv) metabolites with the same ID can be represented with different chemical formula or charge (e.g. “<u>fmn_c</u>“). Therefore, the integration of a BiGG-based model with reactions coming from another BiGG-based model can lead to the introduction of unbalanced reactions or, even worse, stoichiometric inconsistencies [57]. Gempipe tries to circumvent this issue by enabling users to superimpose particular metabolite charges and formulas or reaction balances during the reference expansion phase.

Another limitation depending on BiGG [32] is the limited representation of bacterial diversity, mainly in terms of genes. Indeed, in its current version (v1.6), BiGG contains as little as 108 GSMMs, of which 88 are prokaryotic, of which only 22 do not belong to the *Escherichia* or *Shigella* genera. The set of BiGG genes, used by Gempipe and CarveMe [5], is therefore biased on model species and does not cover much bacterial diversity, potentially leading to missed genes (and thus reactions) during the reference-free reconstruction phase. Gempipe tries to limit this problem by comparing the eggNOG-mapper [58] functional annotation of clusters’ representative sequences (see **Supplementary Information 1.1.2**).

Further development on Gempipe will benefit from the introduction of new universes, such as one for yeasts [59], and from new API functions for the analysis of the deck of strain-specific GSMMs created.

In conclusion, Gempipe will facilitate metabolic biodiversity studies for a wide range of bacterial species, including those not having a dedicated pan-GSMM, which are currently the large majority.

## DATA AVAILABILITY

Gempipe can be easily installed using the dedicated conda package: https://anaconda.org/bioconda/gempipe. Its source code is freely available on GitHub: https://github.com/lazzarigioele/gempipe. Comprehensive documentation for the command-line programs and the API is available on ReadTheDocs, where *ad hoc* tutorials are also included: https://gempipe.readthedocs.io/en/latest/. Code to reproduce validation and case study are available on a separate GitHub repository: https://github.com/lazzarigioele/paper_gempipe. The source code of Cocoremover is available on GitHub: https://github.com/lazzarigioele/cocoremover. All the code used in this paper is also available in Zenodo: https://doi.org/10.5281/zenodo.15544430.

## ACKNOWLEDGEMENTS

/

## AUTHOR CONTRIBUTIONS

**Gioele Lazzari:** Conceptualization, Data curation, Formal analysis, Investigation, Methodology, Software, Validation, Writing – original draft.

**Giovanna E. Felis:** Conceptualization, Data curation, Investigation, Resources, Writing – review & editing.

**Elisa Salvetti:** Data curation, Investigation, Writing – review & editing.

**Matteo Calgaro:** Writing – review & editing.

**Francesca Di Cesare:** Writing – review & editing.

**Bas Teusink:** Supervision, Writing – review & editing.

**Nicola Vitulo:** Conceptualization, Resources, Supervision, Writing – review & editing.

## REFERENCES

1. Domingo-Sananes MR, McInerney JO. Mechanisms That Shape Microbial Pangenomes. Trends in Microbiology 2021;29:493–503.

2. Li W, Wu Q, Kwok L et al. Population and functional genomics of lactic acid bacteria, an important group of food microorganism: Current knowledge, challenges, and perspectives. Food Frontiers 2024;5:3–23.

3. O’Brien EJ, Monk JM, Palsson BO. Using Genome-scale Models to Predict Biological Capabilities. Cell 2015;161:971–87.

4. Thiele I, Palsson BØ. A protocol for generating a high-quality genome-scale metabolic reconstruction. Nat Protoc 2010;5:93–121.

5. Machado D, Andrejev S, Tramontano M et al. Fast automated reconstruction of genome-scale metabolic models for microbial species and communities. Nucleic Acids Research 2018;46:7542–53.

6. Henry CS, DeJongh M, Best AA et al. High-throughput generation, optimization and analysis of genome-scale metabolic models. Nat Biotechnol 2010;28:977–82.

7. Zimmermann J, Kaleta C, Waschina S. gapseq: informed prediction of bacterial metabolic pathways and reconstruction of accurate metabolic models. Genome Biol 2021;22:81.

8. Wang H, Marcišauskas S, Sánchez BJ et al. RAVEN 2.0: A versatile toolbox for metabolic network reconstruction and a case study on Streptomyces coelicolor. Ouzounis CA (ed.). PLoS Comput Biol 2018;14:e1006541.

9. Capela J, Lagoa D, Rodrigues R et al. merlin, an improved framework for the reconstruction of high-quality genome-scale metabolic models. Nucleic Acids Research 2022;50:6052–66.

10. Mendoza SN, Olivier BG, Molenaar D et al. A systematic assessment of current genome-scale metabolic reconstruction tools. Genome Biol 2019;20:158.

11. Monk JM, Charusanti P, Aziz RK et al. Genome-scale metabolic reconstructions of multiple Escherichia coli strains highlight strain-specific adaptations to nutritional environments. Proc Natl Acad Sci USA 2013;110:20338–43.

12. Seif Y, Kavvas E, Lachance J-C et al. Genome-scale metabolic reconstructions of multiple Salmonella strains reveal serovar-specific metabolic traits. Nat Commun 2018;9:3771.

13. Bosi E, Monk JM, Aziz RK et al. Comparative genome-scale modelling of Staphylococcus aureus strains identifies strain-specific metabolic capabilities linked to pathogenicity. Proc Natl Acad Sci USA 2016;113, DOI: 10.1073/pnas.1523199113.

14. Nogales J, Mueller J, Gudmundsson S et al. High-quality genome-scale metabolic modelling of Pseudomonas putida highlights its broad metabolic capabilities. Environ Microbiol 2020;22:255–69.

15. Norsigian CJ, Kavvas E, Seif Y et al. iCN718, an Updated and Improved Genome-Scale Metabolic Network Reconstruction of Acinetobacter baumannii AYE. Front Genet 2018;9:121.

16. Hawkey J, Vezina B, Monk JM et al. A curated collection of Klebsiella metabolic models reveals variable substrate usage and gene essentiality. Genome Res 2022:genome;gr.276289.121v2.

17. Blázquez B, San León D, Rojas A et al. New Insights on Metabolic Features of Bacillus subtilis Based on Multistrain Genome-Scale Metabolic Modeling. IJMS 2023;24:7091.

18. Neal M, Brakewood W, Betenbaugh M et al. Pan-genome-scale metabolic modeling of Bacillus subtilis reveals functionally distinct groups. Faust K (ed.). mSystems 2024:e00923–24.

19. Norsigian CJ, Fang X, Seif Y et al. A workflow for generating multi-strain genome-scale metabolic models of prokaryotes. Nat Protoc 2020;15:1–14.

20. Camacho C, Coulouris G, Avagyan V et al. BLAST+: architecture and applications. BMC Bioinformatics 2009;10:421.

21. Vezina B, Watts SC, Hawkey J et al. Bactabolize: A Tool for High-Throughput Generation of Bacterial Strain-Specific Metabolic Models. elife, 2023.

22. Liao Y-C, Huang T-W, Chen F-C et al. An Experimentally Validated Genome-Scale Metabolic Reconstruction of Klebsiella pneumoniae MGH 78578, i YL1228. J Bacteriol 2011;193:1710–7.

23. Cooper HB, Vezina B, Hawkey J et al. A validated pangenome-scale metabolic model for the Klebsiella pneumoniae species complex. Microbial Genomics 2024;10, DOI: 10.1099/mgen.0.001206.

24. Monk JM. Genome-scale metabolic network reconstructions of diverse Escherichia strains reveal strain-specific adaptations. Phil Trans R Soc B 2022;377:20210236.

25. Hyatt D, Chen G-L, LoCascio PF et al. Prodigal: prokaryotic gene recognition and translation initiation site identification. BMC Bioinformatics 2010;11:119.

26. Seemann T. Prokka: rapid prokaryotic genome annotation. Bioinformatics 2014;30:2068–9.

27. Manni M, Berkeley MR, Seppey M et al. BUSCO Update: Novel and Streamlined Workflows along with Broader and Deeper Phylogenetic Coverage for Scoring of Eukaryotic, Prokaryotic, and Viral Genomes. Kelley J (ed.). Molecular Biology and Evolution 2021;38:4647–54.

28. Shen W, Le S, Li Y et al. SeqKit: A Cross-Platform and Ultrafast Toolkit for FASTA/Q File Manipulation. PLOS ONE 2016;11:e0163962.

29. Rajput A, Chauhan SM, Mohite OS et al. Pangenome analysis reveals the genetic basis for taxonomic classification of the Lactobacillaceae family. Food Microbiology 2023;115:104334.

30. Fu L, Niu B, Zhu Z et al. CD-HIT: accelerated for clustering the next-generation sequencing data. Bioinformatics 2012;28:3150–2.

31. Dimonaco NJ, Aubrey W, Kenobi K et al. No one tool to rule them all: prokaryotic gene prediction tool annotations are highly dependent on the organism of study. Marschall T (ed.). Bioinformatics 2022;38:1198–207.

32. Norsigian CJ, Pusarla N, McConn JL et al. BiGG Models 2020: multi-strain genome-scale models and expansion across the phylogenetic tree. Nucleic Acids Research 2019:gkz1054.

33. Feist AM, Palsson BO. The biomass objective function. Current Opinion in Microbiology 2010;13:344–9.

34. Courtot M, Juty N, Knüpfer C et al. Controlled vocabularies and semantics in systems biology. Molecular Systems Biology 2011;7:543.

35. Lieven C, Beber ME, Olivier BG et al. MEMOTE for standardized genome-scale metabolic model testing. Nat Biotechnol 2020;38:272–6.

36. Moretti S, Tran VDT, Mehl F et al. MetaNetX/MNXref: unified namespace for metabolites and biochemical reactions in the context of metabolic models. Nucleic Acids Research 2021;49:D570–4.

37. Ebrahim A, Lerman JA, Palsson BO et al. COBRApy: COnstraints-Based Reconstruction and Analysis for Python. BMC Syst Biol 2013;7:74.

38. Orth JD, Thiele I, Palsson BØ. What is flux balance analysis? Nat Biotechnol 2010;28:245–8.

39. Li F, Cheng CC, Zheng J et al. Limosilactobacillus balticus sp. nov., Limosilactobacillus agrestis sp. nov., Limosilactobacillus albertensis sp. nov., Limosilactobacillus rudii sp. nov. and Limosilactobacillus fastidiosus sp. nov., five novel Limosilactobacillus species isolated from the vertebrate gastrointestinal tract, and proposal of six subspecies of Limosilactobacillus reuteri adapted to the gastrointestinal tract of specific vertebrate hosts. International Journal of Systematic and Evolutionary Microbiology 2021;71, DOI: 10.1099/ijsem.0.004644.

40. Kristjansdottir T, Bosma EF, Branco dos Santos F et al. A metabolic reconstruction of Lactobacillus reuteri JCM 1112 and analysis of its potential as a cell factory. Microb Cell Fact 2019;18:186.

41. Lee J-Y, Han GG, Choi J et al. Pan-Genomic Approaches in Lactobacillus reuteri as a Porcine Probiotic: Investigation of Host Adaptation and Antipathogenic Activity. Microb Ecol 2017;74:709–21.

42. Abuqwider J, Altamimi M, Mauriello G. Limosilactobacillus reuteri in Health and Disease. Microorganisms 2022;10:522.

43. Yu Z, Chen J, Liu Y et al. The role of potential probiotic strains Lactobacillus reuteri in various intestinal diseases: New roles for an old player. Front Microbiol 2023;14, DOI: 10.3389/fmicb.2023.1095555.

44. Mu Q, Tavella VJ, Luo XM. Role of Lactobacillus reuteri in Human Health and Diseases. Front Microbiol 2018;9, DOI: 10.3389/fmicb.2018.00757.

45. Walter J, Britton RA, Roos S. Host-microbial symbiosis in the vertebrate gastrointestinal tract and the Lactobacillus reuteri paradigm. Proc Natl Acad Sci USA 2011;108:4645–52.

46. Frese SA, Benson AK, Tannock GW et al. The Evolution of Host Specialization in the Vertebrate Gut Symbiont Lactobacillus reuteri. Guttman DS (ed.). PLoS Genet 2011;7:e1001314.

47. Wilson CM, Loach D, Lawley B et al. Lactobacillus reuteri 100-23 Modulates Urea Hydrolysis in the Murine Stomach. Macfarlane GT (ed.). Appl Environ Microbiol 2014;80:6104–13.

48. Bujdoš D, Walter J, O’Toole PW. aurora: a machine learning gwas tool for analyzing microbial habitat adaptation. Genome Biol 2025;26:66.

49. Baroukh C, Cottret L, Pires E et al. Insights into the metabolic specificities of pathogenic strains from the Ralstonia solanacearum species complex. Moleleki L (ed.). mSystems 2023:e00083–23.

50. Notebaart RA, van Enckevort FH, Francke C et al. Accelerating the reconstruction of genome-scale metabolic networks. BMC Bioinformatics 2006;7:296.

51. Battjes J, Melkonian C, Mendoza SN et al. Ethanol-lactate transition of Lachancea thermotolerans is linked to nitrogen metabolism. Food Microbiology 2023;110:104167.

52. De Bernardini N, Zampieri G, Campanaro S et al. pan-Draft: automated reconstruction of species-representative metabolic models from multiple genomes. Genome Biol 2024;25:280.

53. King ZA, Lu J, Dräger A et al. BiGG Models: A platform for integrating, standardizing and sharing genome-scale models. Nucleic Acids Res 2016;44:D515–22.

54. King ZA, Dräger A, Ebrahim A et al. Escher: A Web Application for Building, Sharing, and Embedding Data-Rich Visualizations of Biological Pathways. Gardner PP (ed.). PLoS Comput Biol 2015;11:e1004321.

55. Rowe E, Palsson BO, King ZA. Escher-FBA: a web application for interactive flux balance analysis. BMC Systems Biology 2018;12:84.

56. Seaver SMD, Liu F, Zhang Q et al. The ModelSEED Biochemistry Database for the integration of metabolic annotations and the reconstruction, comparison and analysis of metabolic models for plants, fungi and microbes. Nucleic Acids Research 2021;49:D575–88.

57. Gevorgyan A, Poolman MG, Fell DA. Detection of stoichiometric inconsistencies in biomolecular models. Bioinformatics 2008;24:2245–51.

58. Cantalapiedra CP, Hernández-Plaza A, Letunic I et al. eggNOG-mapper v2: Functional Annotation, Orthology Assignments, and Domain Prediction at the Metagenomic Scale. Tamura K (ed.). Molecular Biology and Evolution 2021;38:5825–9.

59. Lu H, Kerkhoven EJ, Nielsen J. A Pan-Draft Metabolic Model Reflects Evolutionary Diversity across 332 Yeast Species. Biomolecules 2022;12:1632.

