## Supplementary Figures for "Gempipe: a tool for drafting, curating and analyzing pan and multi-strain genome-scale metabolic models"

### MANUSCRIPT TITLE

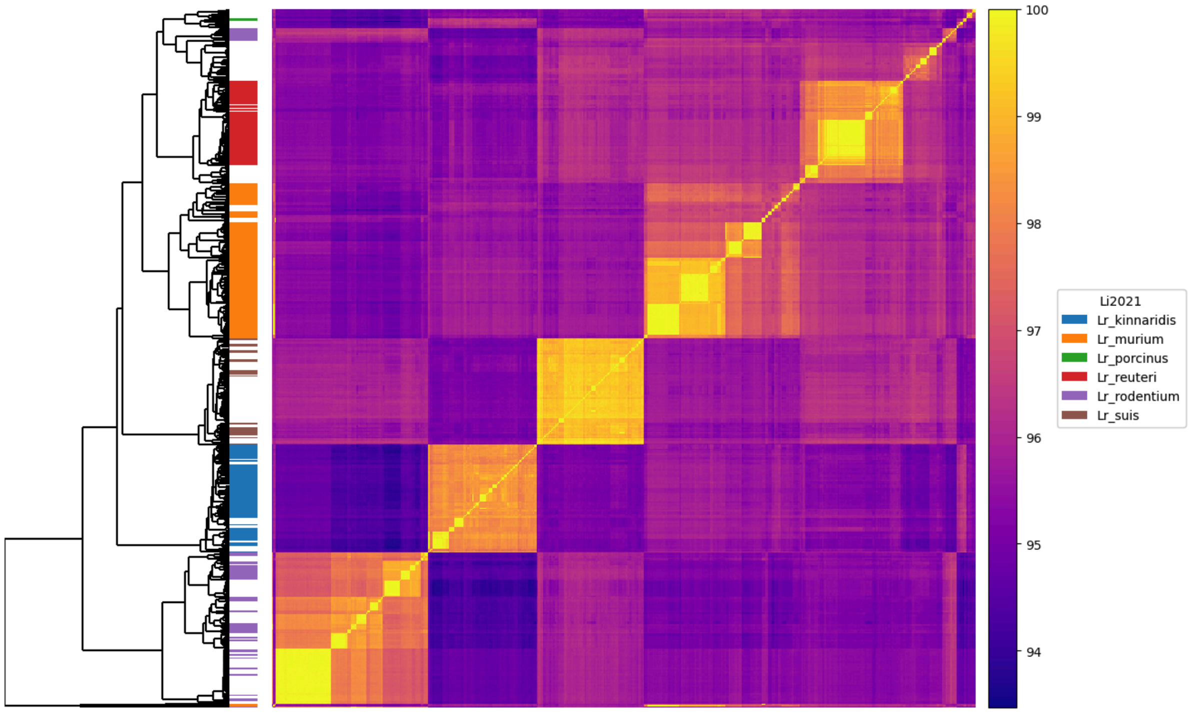


**Figure S1.** ANI matrix for 545 taxonomy- and quality-filtered *Lb* genomes. Genomes were assigned to subspecies by applying ANI thresholds [1] with respect to subspecies type-strains. During this process, not all genomes did receive a subspecies assignment.

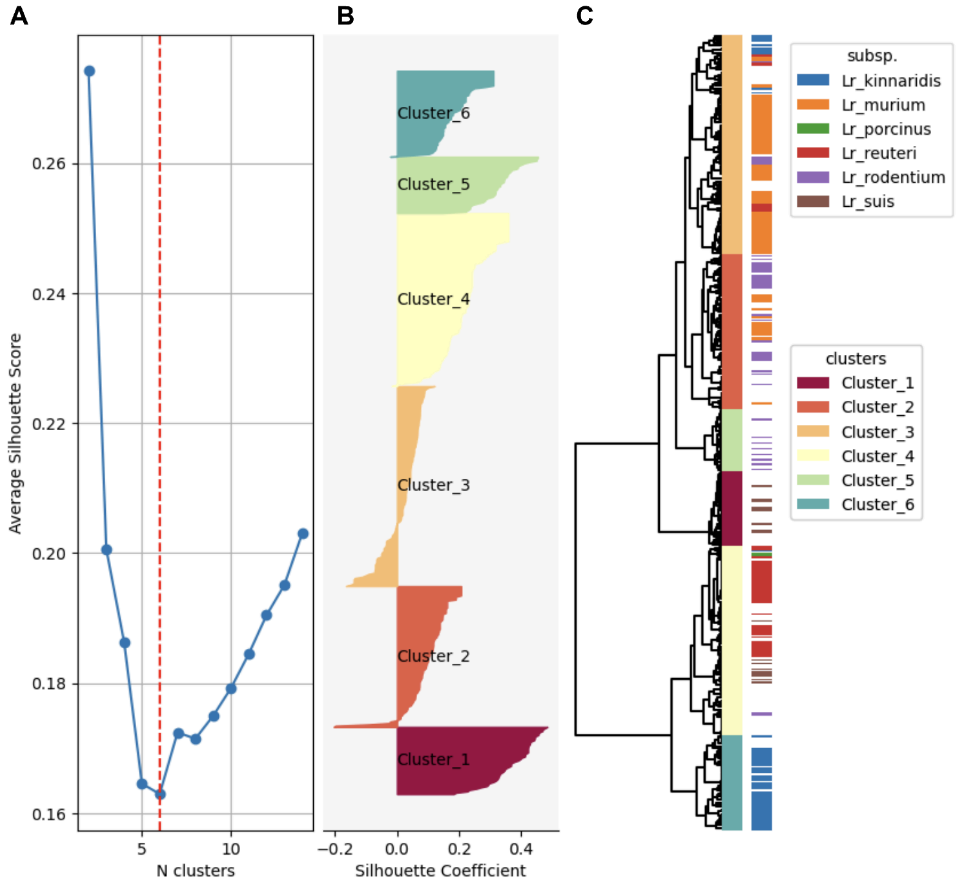


**Figure S2.** Phylometabolic tree and relative clustering of strains without removing the B12 biosynthetic pathway from the dataset (single plot made by the *silhouette_analysis* function from the Gempipe API). **A)** 2 to 14 clusters of metabolically coherent strains were considered for extraction. At first, the average silhouette score suggested the presence of 2 clusters. Indeed, the biosynthetic reactions for vitamin B12 were driving the clusterization of strains into two main groups: producers and non-produces of B12. **B)** 6 clusters of metabolically coherent strains were extracted, according to the number of described subspecies [1]. The silhouette coefficient of individual strains is reported. **C)** Phylometabolic tree built using presence/absence data for reactions, auxotrophies, and growth on alternative C substrates. The subspecies attribute obtained after applying ANI thresholds [1] is reported. The presence/absence of the B12 biosynthetic pathway is separating the strains into 2 main clusters.

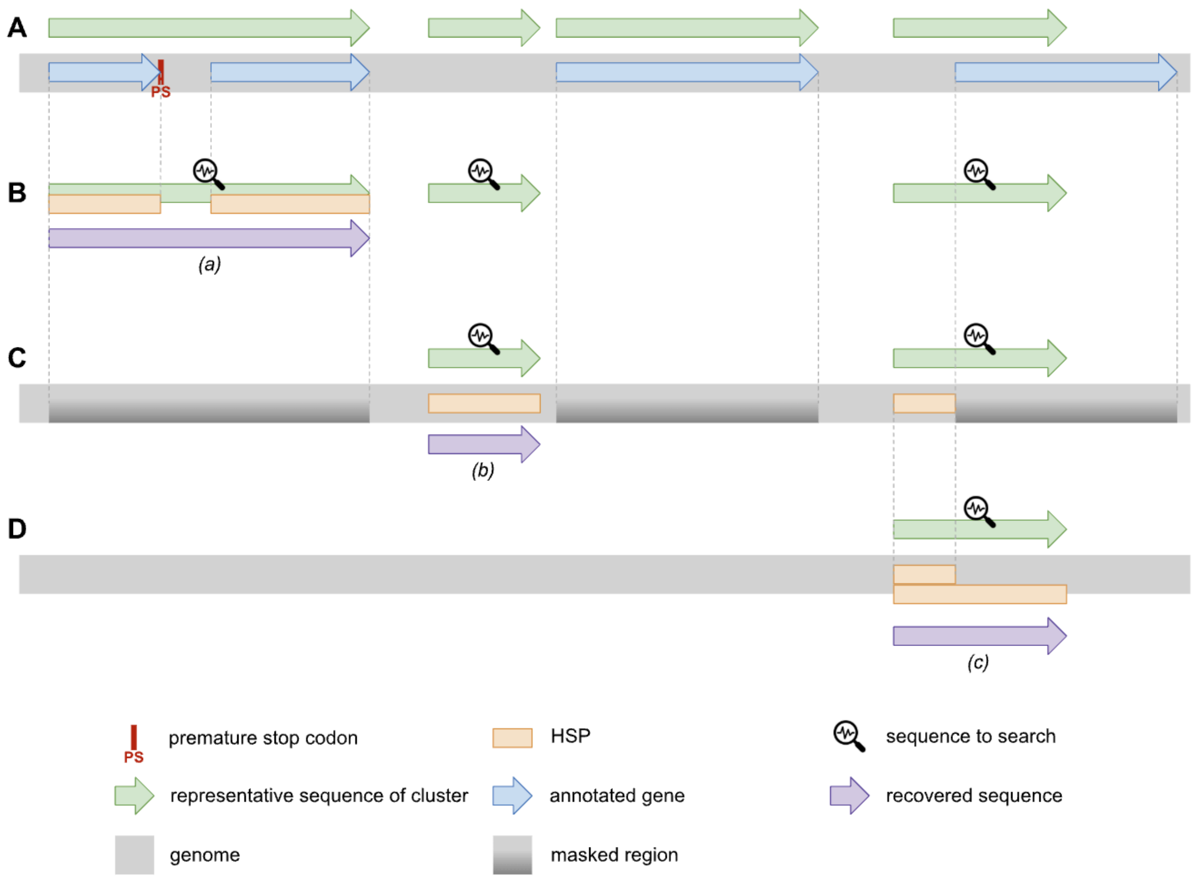


**Figure S3.** General overview of the gene recovery. **A)** Initial gene annotation does not provide strain-specific members for three gene clusters. Sequence *(a)* is broken in two pieces due to the presence of a premature stop codon. Sequence *(b)* falls into a genomic region overlooked by the caller. Sequence *(c)* partially overlaps with an annotated sequence. **B)** First step of gene recovery. Three representative sequences of gene clusters are searched. Annotated genes are aligned on the representative sequences. Two HSPs provide sufficient coverage for a missing sequence, which is recovered. **C)** Second step. Two representative sequences are searched on the masked genome where annotated genes are masked, including those recovered in the previous step. An HSP provides sufficient coverage for a missing sequence, which is recovered. **D)** Third step. One representative sequence is searched on the genome, trying to extend previous HSPs interrupted by masked regions. The extension of a previous HSP provides sufficient coverage for a missing sequence, which is recovered.

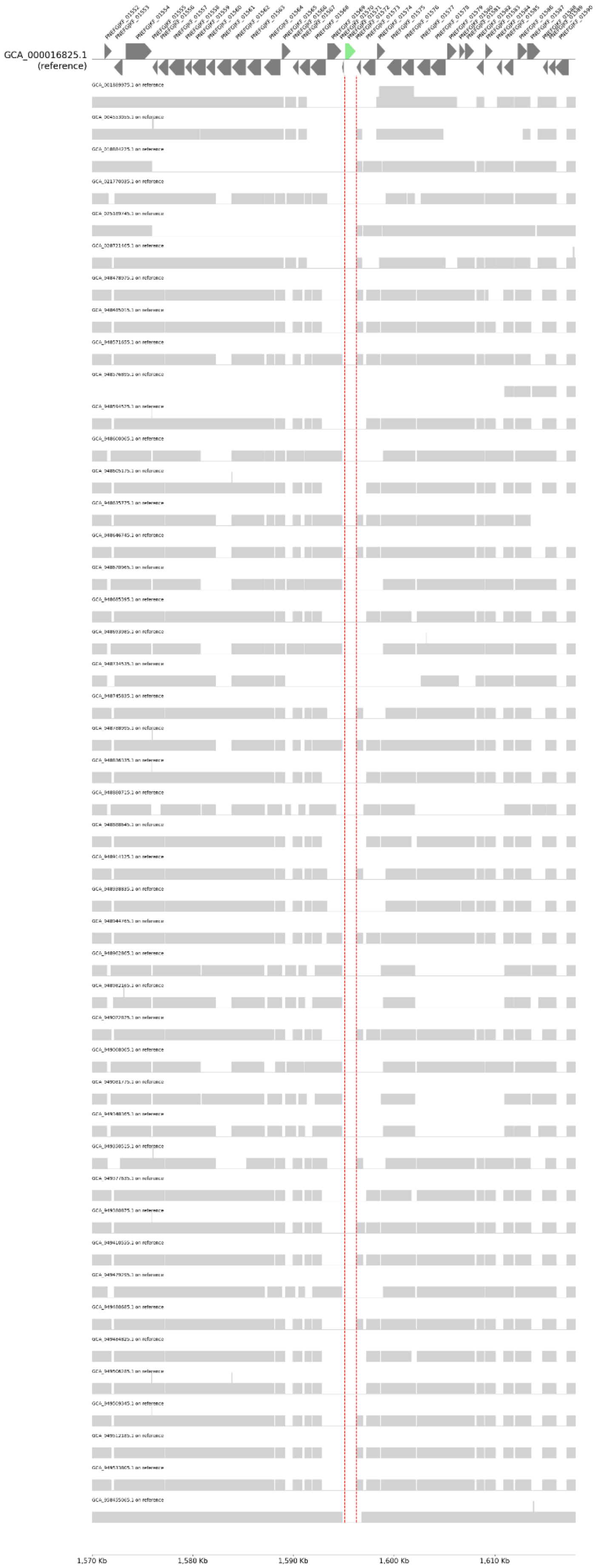
GCA_000016825.1
(reference)
1,570 Kb
1,580 Kb
1,590 Kb
1,600 Kb
1,610 Kb

**Figure S4.** Assembly-on-assembly alignment using the type strain of the species (GCA_000016825.1) as reference. The genomic region displayed is centered on PNEFGJKF_01571 (green arrow), reference gene coding the aldose 1-epimerase (GALM), enzyme used in the first step of the beta-D-galactose catabolism in this species, after the substrate is imported [2]. GALM is missing from ~96% of the 45 strains predicted as unable to catabolize galactose. The interested region is highlighted with dashed red lines.

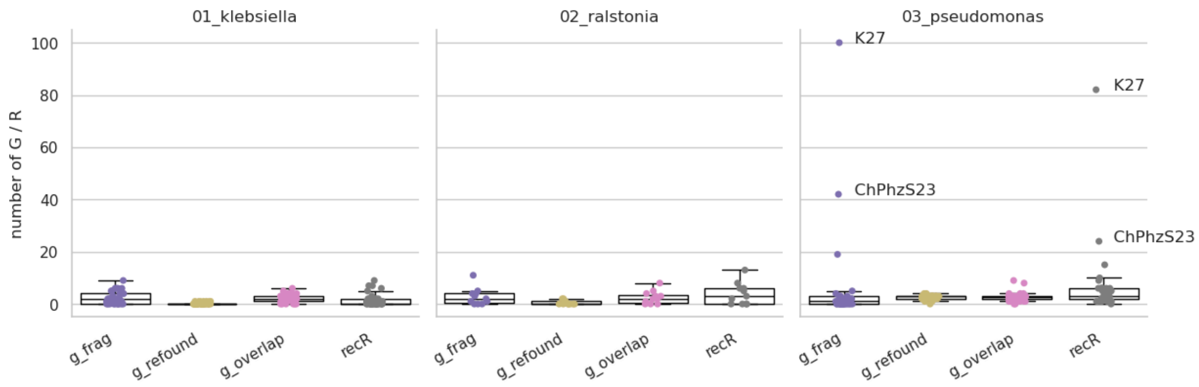


**Figure S5.** Effects of gene recovery. "g_frag", "g_refound" and "g_overlap" are the genes included in strain-specific GSMMs which have been recovered during the first, second, and third step of gene recovery, respectively. "rR" is the number of reactions included in the strain-specific GSMMs consequently to the entire gene recovery procedure.

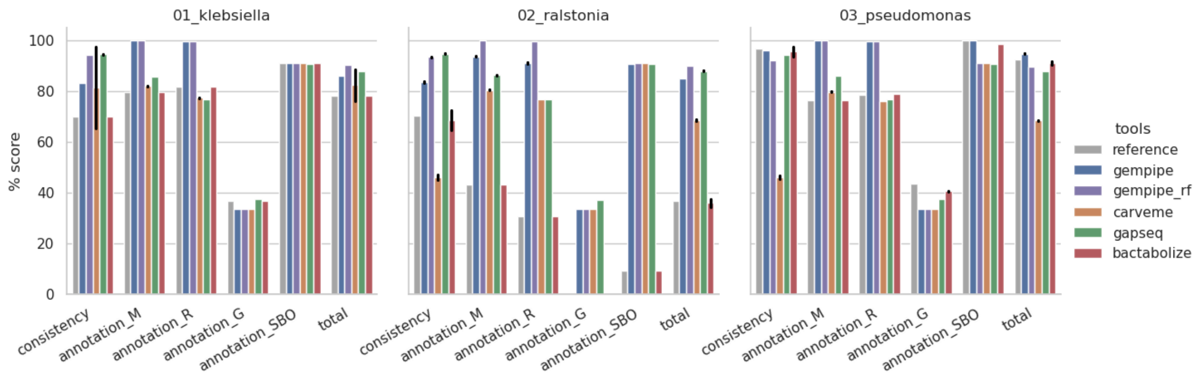


**Figure S6.** Comparison of the MEMOTE metrics and the MEMOTE total score. Bar height corresponds to the mean between strains, while error bars represent standard deviations. "gempipe_rf" indicates Gempipe ran without reference.

### REFERENCES


1. Li F, Cheng CC, Zheng J *et al.* Limosilactobacillus balticus sp. nov., Limosilactobacillus agrestis sp. nov., Limosilactobacillus albertensis sp. nov., Limosilactobacillus rudii sp. nov. and Limosilactobacillus fastidiosus sp. nov., five novel Limosilactobacillus species isolated from the vertebrate gastrointestinal tract, and proposal of six subspecies of Limosilactobacillus reuteri adapted to the gastrointestinal tract of specific vertebrate hosts. *International Journal of Systematic and Evolutionary Microbiology* 2021;**71**, DOI: 10.1099/ijsem.0.004644.

2. Zhao X, Gänzle MG. Genetic and phenotypic analysis of carbohydrate metabolism and transport in Lactobacillus reuteri. *International Journal of Food Microbiology* 2018;**272**:12–21.
