## Supplementary Information for "Gempipe: a tool for drafting, curating and analyzing pan and multi-strain genome-scale metabolic models"

#### MANUSCRIPT TITLE

### 1 SUPPLEMENTARY METHODS

#### 1.1 Implementation

##### *1.1.1 Gene recovery*

When Gempipe starts from genomes, a three-steps gene recovery is applied (**Supplementary Figure S3A**). The steps are based on a BLAST v2.12.0+ [1] alignment where high-scoring segment pairs (HSPs) are filtered with identity ≥ 90%, query coverage ≥ 70%, and e-value ≤ 1e-5, then sorted by ascending e-value unless otherwise stated.

The first step recovers proteins broken into pieces (**Supplementary Figure S3B**) due to eventual issues in genome sequencing or assembling. For each strain, the proteome is aligned with blastp on the representative sequences of clusters. If multiple HSPs share the same query and subject IDs, only the highest-scoring is retained. Prokka [2] names the annotated genes using a progressive number according to the gene order; therefore pairs of sequences named with "n" and "n+1" are searched among HSPs, and the following metrics a calculated: 1) the overall coverage $C$, which includes eventual gap between the two sequences; 2) eventual overlap $S$ between the two sequences; 3) the length $L_1$ and $L_2$ of the sequence pair; 4) the relative frequency $F_1$ and $F_2$ of the starting cluster of each sequence, indicating how many strains in the PAM contain sequences belonging to the cluster; 5) the relative frequency $F_N$ of the new cluster for which the pair members have been grouped. $C$, $S$, $L_1$ and $L_2$ are all expressed in % relatively to the length of the subject (representative sequence). For each detected pair, if $C \geq 70\%$, $S \leq 30\%$, $L_1 < 90\%$, $L_2 < 90\%$, $F_N > F_1$ and $F_N > F_2$, then the pair members – taken together – are assumed as a recovered gene, and the PAM is updated accordingly.

The second gene recovery step deals with unperfect gene calling, in particular with genomic regions that may be overlooked by the caller (**Supplementary Figure S3C**). For each strain, all the coding sequences included in the updated PAM are masked from the original genome. A multi-FASTA query file is then generated, including only the representative sequences of clusters that do not contain proteins belonging to the strain under consideration. Queries are then aligned with tblastn [1] over the masked genome and, for each recovered sequence, the length of the translation until the first stop codon is compared to the length of the full translation: if < 95%, the recovered sequence is assumed to have a premature stop codon, so it will be ignored during GSMMs reconstruction. Finally, the PAM is updated with the recovered sequences.

The third gene recovery step deals with overlapping open reading frames (**Supplementary Figure S3D**). For each strain, a query multi-FASTA is created as in the second step, and queries are then aligned with tblastn [1] over the original (non-masked) genome assembly. For each HSP generated in this step, a HSP from the previous step is searched having same query, same subject (contig), but coverage < 70%. If found, there is evidence of a sequence potentially overlapping with another one previously annotated. After the presence of premature stop codons is verified, the PAM is finally updated with the recovered sequences.

*1.1.2 Reference-free draft pan-GSMM generation*

To create the reference-free draft pan-GSMM, three files are copied from the CarveMe v1.5.2 [3] package, namely "bigg_proteins.faa", "bigg_gprs.csv" and the bacterial universal GSMM. The first is a multifasta file containing all the BiGG-derived [4] gene sequences that constitute the CarveMe gene database (hereafter referred to as the BiGG genes). The second is a table listing, for each BiGG gene, all its GSMM-specific protein complexes in which it is involved, together with the relative reaction ID. The third is a semi-curated, BiGG-based, COBRApy-compatible [5] universal GSMM, either for Gram positive or negative bacteria.

A DIAMOND v2.0.15+ [6] database is created with the BiGG genes sequences, and used to align the representative sequences of clusters with *diamond blastp* program using parameters *--ultra-sensitive --top 10*. A copy of the selected CarveMe universe [3] is made, and all the reactions not supported by the alignment are then removed: the result is the reference-free draft pan-GSMM. In this process, the alignment's bit scores are converted into reaction scores using the information contained in "bigg_gprs.csv", similarly to the CarveMe [3] implementation. However, some important modifications have been implemented to better manage the downstream multi-strain reconstructions.

First, the alignment's HSPs are filtered using user-specified parameters: percentage identity (default 30%) and percentage query and subject coverage (default 70%). This enables users to discard low-confidence reactions based on sequence similarity. Next, each BiGG gene is assigned a score and a partial GPR ("isoform GPR"). This GPR lists all CD-HIT [7] clusters that meet the identity and coverage thresholds for the corresponding BiGG gene, connecting these "isoforms" using the boolean operator "OR". This is a fundamental difference compared to CarveMe [3], which by design does not consider protein isoforms but, for each BiGG gene, it retains just the protein (or cluster, in the case of Gempipe) with the highest bit score, ignoring all the others (see the GitHub public issue #180, https://github.com/cdanielmachado/carveme/issues/180). After this step, each BiGG gene is linked to 0 or more CD-HIT clusters, with an empty isoform GPR in case of 0. To associate a score to each BiGG gene, the maximum bit score among the isoforms is used. Like in CarveMe, the BiGG genes describing spontaneous reactions (artifact genes) are associated with a score of 0.

Next, each protein complex in "bigg_gprs.csv" is associated with a dedicated score and a higher-level partial GPR ("protein complex GPR"). In essence, the isoform GPRs of the BiGG genes involved in the complex are joined together using the boolean operator "AND". The protein complex score is computed as the mean of the gene scores, using 0 for those BiGG genes associated with no clusters. It must be noted that each reaction in the BiGG database can appear in several different BiGG models, and each time it can be described with a different protein complex through its GPR. For example, the mannitol transport reaction "MNLpts" is encoded with 2, 3 or 4 subunits joined by AND operators in model iCN900 [8], iYO844 [9] or iNF517 [10], respectively. In Gempipe, a reaction is copied from the selected universe only if at least one of the BiGG's original protein complex definitions is fully satisfied. For example, in the case of “MNLpts”, if just 1 subunit is found, then the reaction will not be included. Specifically, the protein complex GPR is produced only if all the involved BiGG genes have been associated to an isoform GPR (i.e., at least 1 CD-HIT cluster), this way satisfying the protein complex definitions as they were encoded in the original GSMMs deposited in BiGG [4]. In contrast, CarveMe [3] produces a protein complex GPR with as little as 1 aligned gene, even if the complex was originally defined to be composed by 2 or more members (see the GitHub public issue #182, https://github.com/cdanielmachado/carveme/issues/182).

Lastly, protein complex GPRs and scores are converted into reaction GPRs and scores. As in CarveMe [3], every protein complex GPR linked to the same reaction is joined together using the boolean operator “OR”. The reaction score is computed as the maximum score among the alternative protein complexes, and it is normalized by the median computed among reactions having scores > 0. Given the conservative implementation described above, an empty GPR is assigned to universal reactions having no alternative protein complex fully satisfied. Finally, a copy of the selected CarveMe universe [3] is made and universal reactions with empty GPR are removed. The resulting reconstruction is the reference-free draft pan-GSMM.

Bacterial models in BiGG v1.6 [4], which are not part of the *Escherichia* / *Shigella* species complex, are just 22. This means that BiGG genes are scarce and biased towards model organisms, so they hardly represent the known bacterial diversity. To address this issue, representative sequences of clusters are not only aligned with the BiGG genes database but also functionally annotated using eggNOG-mapper v2.1.7+ [11] using options *-m diamond --itype proteins --trans_table 11*. Then, for each gene cluster involved in the reference-free reconstruction, unmodeled equivalent clusters are retrieved by searching for those having the exact same eggNOG-mapper functional annotation. For each additional gene cluster identified, the involved GPRs in the reference-free reconstruction are expanded accordingly using “OR” statements.

##### *1.1.3 Translation of the reference GSMM*

For each strain, a blastp [1] BRH alignment is performed against the reference proteome. Alignments are parsed to obtain the set of 1-to-1 associations. Next, for each reference gene, all strain-specific 1-to-1 associations are collected to generate a list of strain-specific orthologs. Using the PAM, the CD-HIT [7] cluster corresponding to each ortholog is retrieved, resulting in a list of clusters for each reference gene. Finally, the reference GSMM GPRs are updated by replacing each reference gene with all equivalent cluster IDs connected using the boolean operator “OR”. By removing the original genes, a fully translated version of the reference GSMM is produced.

##### *1.1.4 The reference expansion phase*

Metabolic reactions contained in the repository are iterated, excluding the biomass assembly reaction (“Growth”) and all the spontaneous reactions. For each reaction, an exact string match of reaction ID is performed to check if it is already included in the reference. If the reaction is not found in the reference, its presence with a different ID is checked by searching for reactions having the exact same reactants and metabolites (in terms of IDs), protons excluded, regardless of reversibility or upper and lower bounds. If an equivalent reaction is found in the reference GSMM, a check is made to verify whether it is linked to the same set of cluster IDs. The reference GPR is then updated by taking into account eventual missing cluster IDs.

If an equivalent reaction is not found in the reference GSMM, then the reaction from the repository is assumed as new and it is transferred together with its GPR, bounds and eventual new metabolites. If the new reaction uses metabolites already available in the reference GSMM, then their charge and chemical formula are maintained as defined in the reference. However, the same metabolite (same ID) could be present in different BiGG-based models with different chemical formula and / or charge, and this can lead to the import of unbalanced reactions or, in the worst case, to the violation of the stoichiometric consistency [12]. To avoid such issues, users can provide an optional text file in input, containing a set of rules to be applied during the import of new reactions from the reference-free reconstruction. This enables users to eventually (a) superimpose chemical formula and charge to metabolites being transferred; (b) superimpose a stoichiometric balance to reactions being transferred; (c) exclude reactions from the transfer.

###### *1.1.5 Reannotation of draft pan-GSMM's contents*

To annotate metabolites and reactions, a set of pre-made mapping files is used, mainly derived from MetaNetX v4.4 [13]. Briefly, MetaNetX was downloaded and parsed to obtain the correspondence between BiGG IDs and IDs of several other databases. Regarding metabolites, MetaNetX mappings comprised KEGG (Compound, Drug, and Glycan) [14], MetaCyc [15], HMDB [16], ModelSEED [17], Chebi [18], SABIO-RK [19], LIPID MAPS [20], enviPath [21], Reactome [22], Rhea [23], SwissLipids [24] and BiGG [25] itself. Structural annotations like InChI [26], InChIKey [26], and SMILES [27] were also included. To include the mappings for PubChem [28], the official translations between InChIKey and PubChem IDs were downloaded from the PubChem server (https://ftp.ncbi.nlm.nih.gov/pubchem/RDF/inchikey/pc_inchikey2compound_*.ttl.gz) and parsed. Regarding reactions, MetaNetX mappings comprised KEGG, MetaCyc, ModelSEED, Rhea, SABIO-RK, and BiGG itself, plus the relative EC-codes. To include the mappings for Reactome, the official translations between Rhea and Reactome IDs were downloaded from the Expasy [29] server (https://ftp.expasy.org/databases/rhea/tsv/rhea2reactome.tsv) and parsed. Lastly, to include the mappings for Brenda [30], the official translations were downloaded from the official Brenda website (https://bkms.brenda-enzymes.org/download/Reactions_BKMS.tar.gz) and parsed.

For the annotation of genes, a different approach is used. Gene annotations are retrieved by "gempipe recon" when genbank proteome files are given in input. Genbanks are parsed extracting MIRIAM-compliant [31] gene annotations including "refseq", "ncbiprotein", "ncbigene", "kegg" and "uniprot", which are then applied during the generation of strain-specific GSMMs with "gempipe derive".

Regarding SBO terms [32], the following elements are annotated: genes (SBO:0000243), exchange reactions (SBO:0000627), sink reactions (SBO:0000632), demand reactions

(SBO:0000628), biomass reactions (SBO:0000629), transport reactions (SBO:0000655), purely metabolic reactions (SBO:0000176), and metabolites (SBO:0000247).

###### *1.1.6 Functions for manual curation*

In addition to the command-line programs, Gempipe provides an API which comprises functions for speeding up certain aspects of the manual curation of a GSMM (adding up with those from the community effort MEMOTE [33]). Implemented functions are mainly based on COBRApy v0.29+ [5] and some examples are reported below, grouped by topic.

(i) Sanity check and formatting: identification of energy-generating cycles (EGCs) [34,35]; overview of artifact atoms in metabolites chemical formula (i.e. "X" groups, "R" groups, etc); identification of constrained metabolic reactions. (ii) Network topology: gap-filling for the production of a specified metabolite; biosynthesis verification for metabolites through demand reactions; biosynthesis verification of reactants of a given reaction (e.g. useful to detect blocked biomass precursors); quick import of reactions from a repository GSMM; quick definition of new metabolites and reactions. (iii) Nutritive inputs: simplified configuration of the exchange reactions to represent growth media; sensitivity analysis (eventually scaled) [36] to reveal missing or limiting nutrients.

The API works with any GSMM loaded in COBRAPy [5], but becomes especially useful for the curation of draft pan-GSMMs produced by "gempipe recon", thanks to dedicated functions. For example, the PAM can be quickly queried to retrieve modeled and unmodeled gene clusters based on eggNOG-mapper's functional annotations [11]. Moreover, during the curation of the pan-GSMM, strain-specific GSMMs can be quickly previewed and used to simulate the Biolog® Phenotype MicroArray™ (PM) screening system. This is useful for assessing the effectiveness of the manual curation before starting to derive strain-specific GSMMs with "gempipe derive". Further details and tutorials are available on the Gempipe documentation.

###### *1.1.7 Skipping the manual curation: "gempipe autopilot"*

"gempipe autopilot" creates strain-specific GSMMs directly from genomes or proteomes, skipping the manual curation phase. This is done by internally running "gempipe recon" and "gempipe derive", automatically gap-filling the draft pan-GSMM for biomass production.

The gap-filling is based on an expanded universal GSMM that integrates reactions from the draft pan-GSMM, as well as non-modeled reactions from the selected CarveMe universe and, if provided, the reference GSMM. If a reaction or metabolite appears with identical ID in both the reference GSMM and the CarveMe universe, the version from the reference is chosen.

A prioritized gap-filling is performed using the COBRApy built-in method [5] to ensure biomass production, providing a penalty for each reaction in the expanded universal GSMM. With this approach, missing reactions are added to the draft pan-GSMM favoring those with a better alignment score. The penalty is 0 for reactions already in the draft pan-GSMM, while for others it is calculated as as $1 / (1 + s)$, where $s$ is the normalized reaction score computed using the same method described above for the reference-free draft pan-GSMM reconstruction but with some exceptions. In particular, the alignment HSPs are filtered with relaxed thresholds (identity ≥ 10%, coverage ≥ 40%), allowing more reactions to receive normalized scores beyond those in the draft pan-GSMM. Reactions without genetic support are assigned a score of 0.

User-provided growth media are iteratively applied, with gap-filling performed for each medium. Once the draft pan-GSMM supports biomass production on all media, it is used with the PAM to generate strain-specific GSMMs by running "gempipe derive".

##### *1.1.8 Strain-specific simulations*

Once strain-specific GSMMs are created, *in silico* simulations of the Biolog® Phenotype MicroArray™ (PM) screening system can be performed. Briefly, for each Biolog® PM plate, the substrate present in each well was collected and mapped to the corresponding BiGG [25] exchange reaction. Once the starting sources for C, N, P, and S are defined – by default glucose, ammonia, phosphate, and sulfate, respectively – wells of each plate are iterated. The exchange reaction for the starting C, N, P, or S source is closed (lower bound = 0), depending on how the substrate is categorized in each well (e.g., the L-glutamic acid is tested as a C source in PM01 well B12, but also as a N source in PM03 well A12). Growth is then simulated using flux-balance analysis (FBA) and the objective value ($O_0$) is recorded. Next, the exchange reaction corresponding to the well is opened (unconstrained input) and FBA is performed again, recording the new objective value ($O_1$). If $O_1 \geq O_0 + 0.001$, and the solver status is "optimal", then the strain is considered as able to utilize the substrate of that well.

Detection of auxotrophies can also be performed. Briefly, the production of the following *n* metabolites is tested: all the twenty protein-forming amino acids, plus common vitamins / cofactors (biotine, tetrahydrofolate, lipoate, pantothenate, pyridoxine, pyridoxamine, pyridoxal, riboflavin, thiamine, nicotinate, para-aminobenzoate, cobalamin, ascorbate). The exchange reaction for a compound is closed (lower bound = 0), while those for the other *n*-1 compounds are opened (unconstrained uptake). Then, FBA is performed: if the solver status is "optimal" and the objective value is > 0.001, then the strain is not considered as auxotroph for the compound [37].

A high-throughput screening of growth-enabling alternative substrates can also be performed. Briefly, all the exchange reactions of a strain-specific GSMM are first classified as C, N, P or S sources, depending on the presence of at least one atom of C, N, P or S, respectively (multiple classifications are possible for the same exchange reaction). Once the starting sources for C, N, P, and S are defined – by default glucose, ammonia, phosphate, and sulfate, respectively – all the exchange reactions for C sources are iterated, and substrate consumption is tested like in Biolog® simulations [37]. The procedure is then repeated for N, P, and S sources.

Strain-specific biosynthetic capabilities can also be checked. Briefly, biomass production is constrained to a user-specified fraction of its maximum. Then, for each metabolite present in the model, a demand reaction is temporarily created and set as objective to maximize. After

FBA, if the solver status is "optimal" and the objective value is > 0.001, then the strain is considered a potential producer of the tested metabolite.

##### *1.1.9 Extraction of clusters from phylometabolic trees*

Clusters of metabolically coherent strains can be extracted from a phylometabolic tree. To find the optimal number of clusters, a silhouette analysis [38] is performed, evaluating the clustering quality by measuring how well each data point fits within its own cluster compared to neighboring clusters. The silhouette coefficient quantifies the cohesion and separation of individual data points, ranging from -1 (misclassified) to +1 (well-clustered), providing insight into the suitability of cluster assignments. The average silhouette score (AVS), calculated across all data points, gives an overall measure of clustering efficacy, helping identify optimal cluster numbers and detect poorly formed clusters. The AVS is computed using the *silhouette_score* function from scikit-learn v1.3.0+ (https://github.com/scikit-learn/scikit-learn); the cluster number that maximizes the AVS is used, unless a user-specified number of clusters is specified. Once metabolic clusters are identified on a phylometabolic tree, the cluster attribute can be compared to other strain-specific attributes such as the environmental niche or the species of origin. To identify characteristic features of clusters, features having a relative frequency ≥ *t* and ≤ 1-*t* in two different clusters are selected, where *t* is a customizable threshold with a default value of 0.9.

#### 1.2 Validation

##### *1.2.1 Datasets collection*

To validate and compare Gempipe for the reconstruction of strain-specific GSMMs, publicly available datasets were searched with the following characteristics: (a) organisms belonging to the same species or species complex; (b) publicly available genome assemblies; (c) strain-specific Biolog® PM screenings for carbon sources; (d) published manually curated, phylogenetically close GSMM to be used as reference, belonging at least to the same genus.

Three datasets were collected: (I) *"Klebsiella"* dataset: 37 strains belonging to the *Klebsiella pneumonie* species complex with associated Biolog® data in aerobiose taken from [39]; (II) *"Ralstonia"* dataset: 11 strains belonging to the *Ralstonia solanacearum* species complex with associated Biolog® data in aerobiose taken from [40] (using the time-normalized dataset based on maximal intensity signal); (III) *"Pseudomonas"* dataset: 36 strains of *Pseudomonas chlororaphis* with associated Biolog® data in aerobiose taken from [41]. Strain names and NCBI [42] genome accessions are reported in **Supplementary Table 4**. Details on the respective reference GSMMs are given below.

##### *1.2.2 Reconstruction of strain-specific GSMMs with the tools*

Strain-specific GSMMs were reconstructed for each dataset with four tools: Gempipe v1.38.1 ("autopilot" program), CarveMe v1.6.2 [3], gapseq v1.4 [43] ("*doall*" program), and Bactabolize v1.0.3 [44] ("draft_model" program), all installed via conda. The reference-free reconstruction mode of Gempipe was also evaluated. During reconstructions and simulations, the linear programming solver was CPLEX v22.1.1 [45].

In order to maximize the comparability, all the tools were run providing as input the same amino acid sequences generated by Prodigal v2.6.3 [46] run through Prokka v1.14.6 [2]. The aminoacidic sequences were specified using the *-M 'prot'* option in gapseq, while they are the default input type in CarveMe. In Bactabolize, multifasta proteomes are not accepted as input, therefore the aminoacidic sequences were first formatted with Biopython v1.80 [47] as Bactabolize-compatible genbank files, then the option *--no_reannotation* was used. In Gempipe, the genome filtering feature was not used, to prevent the exclusion of strains during the comparison between tools. Therefore, Gempipe parameters *--buscoM*, *--ncontigs* and *--N50* were set to 100%, 10000 and 0, respectively.

The reference GSMM and associated coding sequences were specified using the options *-rm/-rp* in Gempipe, *--reference* in CarveMe, and *--ref_model_fp/--ref_proteins_fp/--ref_genes_fp* in Bactabolize. In gapseq, it is not possible to specify a reference GSMM, therefore reconstructions are fully based on the gapseq internal universe. Reconstructions are purely universe-based also in CarveMe, but the *--reference* option increases the reaction scores of those reactions in common with the reference, based on reaction ID exact matching [3]. Moreover, the use of the reference GSMM as an alternative CarveMe universe already showed poor performances [44], so this reconstruction mode was ignored here. The gram-negative universe was specified with *-s* in Gempipe and *-u* in CarveMe, while gapseq relies on a staining autodetection feature.

As reference for the "*Klebsiella"* dataset, the curated model iYL1228 [48], specific for *Klebsiella pneumonie* MGH 78578, was downloaded from BiGG [4] and slightly edited as follows. The biomass equation was edited by removing the strain-specific precursors dTDP-rhamnose ("dtdprmn_c"), UDP-galacturonate ("udpgalur_c") and UDP-galactose ("udpgal_c"), as indicated in [39]. Diffusion reactions involved in the transport of Biolog® substrates were systematically set as reversible [39]. Modeled gene sequences were retrieved from NCBI (CP000647.1).

As reference for the "*Ralstonia"* dataset, the curated model iRP1476 [49], specific for *Ralstonia solanacearum* GMI1000, was downloaded in a COBRApy-compatible version from supplementary materials of [3], and slightly edited as follows. All unconserved metabolites were detected with MEMOTE v0.17.0 [33] and manually corrected. The gene RSp042 was changed to RSp0421 in the "ARBTNLSYN" reaction GPR. To uniform the reconstruction paradigm among models, prefix "G_" was removed from gene IDs; genes whose ID started with "e" (associated to exchange reactions) were removed; genes whose ID started with "d" or "s" (associated to diffusion and spontaneous reactions, respectively) were all converted to an artifact "spontaneous" gene; the gene "NoAssignment" was removed from GPRs. As indicated in [40], cobalamin ("adocbl_c") and sperimidine ("spmd_c") are non-essential for growth, and were thus removed from the biomass assembly. Finally, an uptake for $Ca^{2+}$ was introduced as a proton antiport. Modeled gene sequences were retrieved from NCBI (NC_003295.1, NC_003296.1, AL646052.1, AL646053.1). Modeled genes with no available sequence were removed from the model without removing reactions.

As reference for the "*Pseudomonas*" dataset, the curated model iJN1463 [50], specific for *Pseudomonas putida* KT2440, was downloaded from BiGG [25] and slightly edited as follows. Its unconstrained bound constants were changed from +/- 999999 to +/- 1000, and the objective reaction was set to the preloaded "core" biomass equation described in [50]. Modeled gene sequences were retrieved from NCBI (NC_002947.4). Modeled genes with no available sequence were removed from the model without removing reactions.

The growth media used during gap-filling was specified using the options *--media* in Gempipe, *-g/--mediadb* in CarveMe, *-n* in gapseq, *--media_type/--atmosphere_type* in Bactabolize. Unconstrained exchange of protons, water, oxygen, and $CO_2$ was allowed. In Bactabolize, oxygen and $CO_2$ were not included in the medium definition as the *--atmosphere_type aerobic* option was used. Trace elements were approximated as an unconstrained uptake of $Ca^{2+}$, $Cl^-$, $Co^{2+}$, $Cu^{2+}$, $Fe^{2+}$, $Fe^{3+}$, $K^+$, $Mg^{2+}$, $Mn^{2+}$, $Zn^{2+}$ and $MoO_4^{2-}$. N, P, and S sources during the gap-filling phase were ammonium, phosphate and sulfate, respectively, indicated as unconstrained uptake. C source uptake was constrained to 10 mmol/gDW/h, and was glucose for the "*Klebsiella*" and "*Pseudomonas*" datasets, and L-glutamine for the "*Ralstonia*" dataset. For the "*Pseudomonas*" dataset, an additional uptake of $Na^+$ and $Ni^{2+}$ was necessary to satisfy the growth requirements of iJN1463. As Bactabolize does not include an automatic gap-filling feature, models generated with Bactabolize were automatically gap-filled with COBRApy v0.29.0 [5] using the specified growth medium and the reference GSMM as source of reactions, requiring a minimum growth rate of 0.6 1/h. For the gap-filling in Gempipe, the minimum growth rate was set to 0.9 1/h for the pan-GSMM and 0.6 1/h for the strain-specific GSMMs, using the *--minpanflux* and *--minflux* options, respectively.

To avoid stoichiometric inconsistencies in the GSMMs produced by Gempipe, a blacklist of reactions not to be included during the expansion phase was set using *--mancor*. Specifically, the set was ("ACPpds", "ENTERH", "NFORGLUAH2"), ("ACPpds"), and ("AALDH", "ACPpds", "ASR2", "PROR") to for the "*Klebsiella*", "*Ralstonia*" and "*Pseudomonas*" dataset, respectively. The custom reference artifact gene indicating spontaneous reactions was specified in Gempipe as *-rs KPN_SPONT* and *-rs PP_s0001* for the "*Klebsiella*" and "*Pseudomonas*" datasets, respectively.

##### *1.2.3 Comparison of contents*

Each strain-specific GSMM was loaded into COBRApy [5], and four sets of IDs were extracted: for genes ("G"), reactions ("R"), metabolites ("uM") and exchange reactions ("exr"). Metabolite IDs were extracted without considering their compartment. Moreover, the number of reactions without GPR (orphans) was computed ignoring reactions with either just one metabolite involved (exchanges, sinks, and demands) or with "diffusion" present in the name (reactions marked as spontaneous were not considered). For GSMMs created by gapseq, the IDs of metabolites and reactions were translated from SEED to BiGG namespace using the MetaNetX v4.4 [13] mappings.

Each GSMM was then evaluated through the computation of MEMOTE metrics [33] which include five categories: consistency, metabolite annotations, reaction annotations, gene annotations, and SBO annotations, along with a total score. Metrics were calculated using the MEMOTE API for Python (*memote.suite.api.test_model*). The results replicated the original MEMOTE metrics by applying the formulas described in the original paper [33].

To assess the reference coverage, i.e. the degree of superimposition between the output models and the reference, the intersection between each strain-specific GSMM and its reference was computed in terms of ID sets. To intersect genes, each reference gene ID was first converted to the corresponding set of cluster IDs, then all the associated strain-specific gene IDs were collected by reading the PAM. A reference gene was considered as "covered" if at least one of these equivalent genes was present in the strain-specific GSMM.

To compare the similarity among the reconstruction tools in terms of reactions, a correlation matrix was computed. Each cell of the matrix reported the similarity among two tools, represented by the mean Jaccard index

$$\sum_{i=1}^{n}\left(\frac{R_i^A \cap R_i^B}{R_i^A \cup R_i^B}\right)\frac{1}{n}$$

where $R_i^A$ and $R_i^B$ are the BiGG-based set of reaction IDs of the tool *A* and *B*, respectively, and *n* is the number of strains in the dataset; when a tool is compared against a reference GSMM, then each strain-specific GSMM is compared against a copy of the reference.

##### *1.2.4 Comparison of the simulations*

For all datasets and tools, Biolog® substrate utilization was simulated using the *biolog_preview* function from the Gempipe API, indicating glucose, ammonium, phosphate and sulfate as starting C, N, P, and S sources, respectively. The argument *seed=True* was utilized for gapseq reconstructions. If the exchange reaction for a particular substrate was missing from a GSMM, this was interpreted as inability to catabolize that substrate.

To compare substrate utilization predictions among tools, the following metrics were considered [3,44]:

precision $= \frac{TP}{TP+FP}$

recall $= \frac{TP}{TP+FN}$

specificity $= \frac{TN}{TN+FP}$

accuracy $= \frac{TP+TN}{TP+TN+FP+FN}$

TP, TN, FP and FN are the number of true positive, true negative, false positive and false negative substrates, respectively. The cases of no growth and infeasible solution were distinguished: the first case is when no growth is observed *in silico* while the solver status is "optimal"; the second case is when the linear programming problem defined by the FBA has no valid solution, not even 0, corresponding to the solver status "infeasible". To penalize reconstructions with significant gaps, each of the above metrics was multiplied by the fraction of "optimal" (non-"infeasible") FBAs, where each FBA corresponds to a specific substrate.

#### 1.3 *Limosilactobacillus reuteri* case study

##### *1.3.1 Genome decontamination, filtering and species assignation*

All the genome assemblies associated with taxid 1598, belonging to *Limosilactobacillus reuteri* (*Lr*), were downloaded from NCBI GenBank [42] in their latest version. Assemblies labeled by NCBI as "derived from metagenome" were subjected to decontamination, a process where contigs belonging to species other than *Lr* were removed. Briefly, all amino acidic protein sequences associated to genomes from type-materials were retrieved from NCBI RefSeq [42], associated to the respective species-level NCBI Taxonomy ID, and finally used to create a Diamond [6] reference database. For each input assembly, genes were predicted with Prodigal [46] and aligned on the reference database using the *diamond blastp* program (default options). For each predicted gene, the HSP with the higher bit-score was used to associate the gene with a species. For each contig of the input assembly, if the species associated with the higher number of genes was not *Lr*, then the contig was assumed as contaminant and removed from the assembly. This method, together with the procedure to build an updated reference database, was packaged as a separate tool named Cocoremover, and made publicly available. Cocoremover v0.1.0 was used for the decontamination, building the reference database on 16th May 2025.

Genomes' average nucleotide index (ANI) was computed with FastANI v1.34 [51] and a taxonomy filter was applied by removing all genomes having ANI < 95% with respect to the type strain of the species (GCA_000016825.1). A quality filter was further applied by Gempipe v1.38.1 using parameters *-b lactobacillales_odb10 --buscoM 3% --buscoF 2% --ncontigs 240 --N50 19000* in "gempipe recon".

Remaining genomes were assigned to *Lr* subspecies by using thresholds reported by Li *et al.* [52]. Specifically, each strain was assigned to a subspecies if its ANI was ≥ *t* respect to the type strain of the subspecies, where *t* is 98.1% for "reuteri", 98.2% for "kinnaridis", 99.1% for "porcinus", 96.8% for "murium", 98.7% for "suis", and 96.1% for "rodentium" [52]. During this procedure, several genomes were not assigned to any subspecies. To guarantee a confident subspecies classification, strains assigned to 2 or more subspecies were considered as unclassified.

##### *1.3.2 GSMMs reconstruction and simulations*

The cured model Lreuteri_530, specific for *Lr* JCM1112, was downloaded from [53], and used as reference GSMM in "gempipe recon" after the following edits. Metabolites with no linked compartment were assigned to the compartment indicated in their ID. The growth media components were maintained as indicated in [53], but their respective exchange reaction bounds were qualitatively uniformed to (-5; 1000) with glucose as only exception (-10; 1000); all the other exchange reactions were relaxed to (0; 1000).

A draft pan-GSMM was built using "gempipe recon" with option *-s pos* (gram-positive universe), expanding the reference GSMM with new strain-specific reactions encoded in the filtered genomes. Using the *verify_egc* function from the Gempipe API, an EGC for the production of ATP was detected in the draft pan-GSMM and consequently corrected. Moreover, before using the pan-GSMM as input for "gempipe derive", vitamin B12 (cobalamin, *adeadocbl_c*) was removed from the biomass equation, as its production was proved to be strain-specific [54–56].

The updated pan-GSMM was given in input "gempipe derive" to generate strain-specific GSMMs. For the strain-specific gap-filling, the medium was kept the same described above (*--media*) and the minimum flux through the objective was set to 0.5 (*--minflux*). Moreover, the generation of BFTs for auxotrophies, alternative substrates, and biosynthetic abilities were requested with the dedicated options (*--aux*, *--cnps*, *–biosynth 0.8*, respectively), in addition to the BFT for the strain-specific reaction content (produced by default). Subsequently, the BFT for alternative substrates was filtered to retain only the C sources.

##### *1.3.3 Multi-strain analysis and comparisons*

The 3 BFTs (reactions, auxotrophies, alternative substrates) were given in input to the *silhouette_analysis* function from the Gempipe API, to perform a silhouette analysis, produce a phylometabolic tree and extract metabolic clusters. The same procedure was repeated after removing from the feature set the 24 reactions composing the biosynthetic pathway for vitamin B12 ('GLUTRR', 'G1SAT', 'PPBNGS', 'HMBS', 'UPP3S', 'UPP3MT', 'SRCHCOC', 'COPREC2MT', 'COPREC3MT', 'COPREC4MT', 'COPRECT', 'COPREC6R2', 'COPREC6R2', 'COPREC6MT', 'COPRECI', 'CBIA', 'R05218', 'R05220', 'R05225', 'ADCOBAPS', 'ACBIPGT', 'NNDMBRT', 'ADEADOCBLS', 'ADEADOCBLPP'), as the presence/absence of the underlying operons is not related to host adaptation [56]. The variable feature content associated with the phylometabolic tree was visualized with the *heatmap_multilayer* function from the Gempipe API. Relative frequency of metabolic features in the clusters was computed with the *discriminant_feat* function from the Gempipe API.

Experimental C substrate consumptions for *Lr* were retrieved from Li *et al.* [52] to be compared with *in silico* predictions. If a substrate had no corresponding exchange reaction in the draft pan-GSMM, no consumption was assumed for all the strains. Substrates were grouped into 2 datasets according to Li *et al.* [52]: a "general" dataset, containing the description of the subspecies (1: phenotypic characteristic always present in all the strain of the subspecies; 0 always absent; 0.5: strain-specific characteristic); a "strain-specific" dataset, containing phenotypic characteristic of 12 individual *Lr* strains, 2 strains for each *Lr* subspecies.

The "strain-specific" dataset was compared directly with the predictions obtained from the corresponding GSMM. On the other hand, to compare the "general" dataset with predictions, each subspecies was first assigned to the cluster with the most strains of that subspecies, based on the previous ANI threshold assignment. Then, the relative frequency of metabolic features in each cluster was used for comparison, considering also the contribution of strains that were not initially assigned to that subspecies. To check the deletion of key genes, minimap v2.28 [57,58] with options *-ax asm5* was used to align the genome assembly of a strain on the assembly of the type strain of the species (GCA_000016825.1), used as reference. The library pysam v0.21.0 [59] (https://github.com/pysam-developers/pysam) was used to extract a pileup of the target genomic region, which was then plotted using the library pyGenomeViz v1.6.0 (https://github.com/moshi4/pyGenomeViz).

Finally, the presence of 4 key metabolic features was investigated. Specifically, the potential biosynthesis of vitamin B12 ("adeadocbl_c"), reuterin ("3hppnl_c"), histamine ("hista_c"), together with the potential conversion of urea to ammonium and $CO_2$ ("UREA"), were evaluated. A dedicated plot was made with the *heatmap_multilayer* function from the Gempipe API.

#### 2 SUPPLEMENTARY RESULTS

##### 2.1 Galactose-deficient strains

In *Lr*, galactose enters the cell with a proton symporter (GALt), gets converted to alpha-D-galactose with an aldose 1-epimerase (GALM), then to galactose-1P with a galactokinase (GALK2), then to UDP-galactose with a galactose-1-phosphate uridylyl-transferase (GALT), then to UDP-glucose with an UDP-glucose 4-epimerase (UDPG4E) and finally to glucose-1P with an UDP-glucose pyrophosphorylase (GALU) [60]. The six correspondent reactions were present in the 45 strains with a relative frequency of 0.98, 0.04, 0.98, 1.00, 1.00 and 1.00, respectively. Therefore, the deletion of the aldose 1-epimerase was checked (**Supplementary Figure S4**): while metagenome-derived assemblies might not be complete, a consistent gap in GALM was observed in strains unable to catabolize galactose.

To exclude the possibility that simulated phenotypes deviated from literature because of the inclusion of strains assigned to different subspecies in the same cluster, relative frequencies were computed again on groups of strains strictly belonging to the same subspecies. Even in this case, the relative frequencies differed from literature data. For example, strains strictly classified as *murium* and unable to catabolize galactose increased to ~26%.

##### 2.2 Effects of the gene recovery

As detailed in **Supplementary Methods 1.1**, Gempipe provides a gene recovery feature to cope with possible errors due to sequencing, assembling, or gene prediction. This recovery is divided in three steps executed in order: the first step tries to reconstitute proteins broken in two pieces, the second searches for missing genes along unconsidered genomic regions, while the third deals with overlapping open reading frames [61].

In general, as expected, the entire gene recovery procedure was conservative for all the tested datasets, as the overall number of recovered genes (**Supplementary Table 5**) and reactions (**Supplementary Table 6**) contained in strain-specific GSMMs (**Supplementary Figure S5**) resulted low compared to the total (**Figure 2**), with a median of 5 recovered genes and 2 recovered reactions, regardless the dataset.

However, two evident outliers were found, namely strains K27 and ChPhzS23 from the dataset *Pseudomonas*, both sequenced with PacBio technology [62]. Their corresponding accessions were both flagged on NCBI as *"removed from RefSeq due to many frameshift proteins"*, therefore these genomes should normally be excluded from an analysis. Furthermore, the 100% of the recovered reactions for these two strains were always present in the other strains of the dataset, demonstrating that the gene recovery procedure is effective in improving the metabolism reconstruction even for genomes with low sequence quality.

##### 2.3 MEMOTE metrics evaluation

As an additional comparison between tools, the scores obtained by MEMOTE [33], a widely used tool for assessing the quality of GSMMs, were evaluated (**Supplementary Figure**

**S6**). Five metrics are merged into a total score: stoichiometric consistency and metabolite, reaction, gene, and SBO annotations [32]. Of these, stoichiometric consistency, the condition for which a positive mass can be associated to all metabolites in a GSMM [12], is weighted the most in the total score [33].

In general, all tools achieved high scores in terms of stoichiometric consistency, except for CarveMe, which violated consistency in two datasets, despite CarveMe universes being curated for consistency. Indeed, Gempipe in reference-free mode, which used the same CarveMe universes, scored higher in terms of consistency.

The annotations provided by Bactabolize are completely inherited from the reference, meaning that strain-specific GSMMs cannot score better than the reference itself. Similarly, CarveMe and gapseq are bonded to the annotation of their internal universes. In contrast, Gempipe's re-annotation based on MetaNetX [13] produced consistently higher scoring annotations, particularly for metabolites and reactions.

### 3 SUPPLEMENTARY DISCUSSION

#### 3.1 Insights obtained by modeling *L. reuteri* strains

Gempipe was used to process more than 1000 publicly available *L. reuteri* genome assemblies, grouping strains into metabolically coherent clusters. Although clustering was essentially driven by the presence/absence of metabolic reactions excluding other strain-specific metabolic traits, such as the host-adapted adhesion proteins [63,64], the resulting clusters resembled the subspecies division proposed in literature [52]. However, per-strain prediction of metabolic capabilities revealed that characterizing as few as two strains [52] was probably insufficient to accurately define the general metabolic profile of a subspecies. Moreover, the multi-strain analysis enabled by Gempipe showed that, when analysing a number of strains larger than that analysed for taxonomic purposes, metabolic boundaries between subspecies become more fuzzy, suggesting a metabolic continuum within the species, rather than stable and robust infraspecific separation of strains. Therefore this kind of analysis can push forward and improve taxonomic interpretation of biodiversity, especially within species level, in line with other suggestions [65]. Further, it shows a great potential in strain screening and rational selection, as demonstrated in this study case by focusing on health-related key metabolites including ruterin, vitamin B12 and histamine.

#### REFERENCES

1. Camacho C, Coulouris G, Avagyan V *et al.* BLAST+: architecture and applications. *BMC Bioinformatics* 2009;**10**:421.

2. Seemann T. Prokka: rapid prokaryotic genome annotation. *Bioinformatics* 2014;**30**:2068–9.

3. Machado D, Andrejev S, Tramontano M *et al.* Fast automated reconstruction of genome-scale metabolic models for microbial species and communities. *Nucleic Acids Research* 2018;**46**:7542–53.

4. Norsigian CJ, Pusarla N, McConn JL *et al.* BiGG Models 2020: multi-strain genome-scale models and expansion across the phylogenetic tree. *Nucleic Acids Research* 2019:gkz1054.

5. Ebrahim A, Lerman JA, Palsson BO *et al.* COBRApy: COnstraints-Based Reconstruction and Analysis for Python. *BMC Syst Biol* 2013;**7**:74.

6. Buchfink B, Xie C, Huson DH. Fast and sensitive protein alignment using DIAMOND. *Nat Methods* 2015;**12**:59–60.

7. Fu L, Niu B, Zhu Z *et al.* CD-HIT: accelerated for clustering the next-generation sequencing data. *Bioinformatics* 2012;**28**:3150–2.

8. Norsigian CJ, Danhof HA, Brand CK *et al.* Systems biology analysis of the Clostridioides difficile core-genome contextualizes microenvironmental evolutionary pressures leading to genotypic and phenotypic divergence. *npj Syst Biol Appl* 2020;**6**:31.

9. Oh Y-K, Palsson BO, Park SM *et al.* Genome-scale Reconstruction of Metabolic Network in Bacillus subtilis Based on High-throughput Phenotyping and Gene Essentiality Data. *Journal of Biological Chemistry* 2007;**282**:28791–9.

10. Flahaut NAL, Wiersma A, Van De Bunt B *et al.* Genome-scale metabolic model for Lactococcus lactis MG1363 and its application to the analysis of flavor formation. *Appl Microbiol Biotechnol* 2013;**97**:8729–39.

11. Cantalapiedra CP, Hernández-Plaza A, Letunic I *et al.* eggNOG-mapper v2: Functional Annotation, Orthology Assignments, and Domain Prediction at the Metagenomic Scale. Tamura K (ed.). *Molecular Biology and Evolution* 2021;**38**:5825–9.

12. Gevorgyan A, Poolman MG, Fell DA. Detection of stoichiometric inconsistencies in biomolecular models. *Bioinformatics* 2008;**24**:2245–51.

13. Moretti S, Tran VDT, Mehl F *et al.* MetaNetX/MNXref: unified namespace for metabolites and biochemical reactions in the context of metabolic models. *Nucleic Acids Research* 2021;**49**:D570–4.

14. Kanehisa M. KEGG: Kyoto Encyclopedia of Genes and Genomes. *Nucleic Acids Research* 2000;**28**:27–30.

15. Caspi R, Altman T, Billington R *et al.* The MetaCyc database of metabolic pathways and enzymes and the BioCyc collection of Pathway/Genome Databases. *Nucl Acids Res* 2014;**42**:D459–71.

16. Wishart DS, Guo A, Oler E *et al.* HMDB 5.0: the Human Metabolome Database for 2022. *Nucleic Acids Research* 2022;**50**:D622–31.

17. Seaver SMD, Liu F, Zhang Q *et al.* The ModelSEED Biochemistry Database for the integration of metabolic annotations and the reconstruction, comparison and analysis of metabolic models for plants, fungi and microbes. *Nucleic Acids Research* 2021;**49**:D575–88.

18. Hastings J, Owen G, Dekker A *et al.* ChEBI in 2016: Improved services and an expanding collection of metabolites. *Nucleic Acids Res* 2016;**44**:D1214–9.

19. Wittig U, Kania R, Golebiewski M *et al.* SABIO-RK--database for biochemical reaction kinetics. *Nucleic Acids Research* 2012;**40**:D790–6.

20. Conroy MJ, Andrews RM, Andrews S *et al.* LIPID MAPS: update to databases and tools for the lipidomics community. *Nucleic Acids Research* 2024;**52**:D1677–82.

21. Wicker J, Lorsbach T, Gütlein M *et al.* enviPath – The environmental contaminant biotransformation pathway resource. *Nucleic Acids Res* 2016;**44**:D502–8.

22. Milacic M, Beavers D, Conley P *et al.* The Reactome Pathway Knowledgebase 2024. *Nucleic Acids Research* 2024;**52**:D672–8.

23. Bansal P, Morgat A, Axelsen KB *et al.* Rhea, the reaction knowledgebase in 2022. *Nucleic Acids Research* 2022;**50**:D693–700.

24. Aimo L, Liechti R, Hyka-Nouspikel N *et al.* The SwissLipids knowledgebase for lipid biology. *Bioinformatics* 2015;**31**:2860–6.

25. King ZA, Lu J, Dräger A *et al.* BiGG Models: A platform for integrating, standardizing and sharing genome-scale models. *Nucleic Acids Res* 2016;**44**:D515–22.

26. Heller SR, McNaught A, Pletnev I *et al.* InChI, the IUPAC International Chemical Identifier. *J Cheminform* 2015;**7**:23.

27. Weininger D. SMILES, a chemical language and information system. 1. Introduction to methodology and encoding rules. *J Chem Inf Comput Sci* 1988;**28**:31–6.

28. Kim S, Chen J, Cheng T *et al.* PubChem 2023 update. *Nucleic Acids Research* 2023;**51**:D1373–80.

29. Gasteiger E. ExPASy: the proteomics server for in-depth protein knowledge and analysis. *Nucleic Acids Research* 2003;**31**:3784–8.

30. Chang A, Jeske L, Ulbrich S *et al.* BRENDA, the ELIXIR core data resource in 2021: new developments and updates. *Nucleic Acids Research* 2021;**49**:D498–508.

31. Juty N, Le Novere N, Laibe C. Identifiers.org and MIRIAM Registry: community resources to provide persistent identification. *Nucleic Acids Research* 2012;**40**:D580–6.

32. Courtot M, Juty N, Knüpfer C *et al.* Controlled vocabularies and semantics in systems biology. *Molecular Systems Biology* 2011;**7**:543.

33. Lieven C, Beber ME, Olivier BG *et al.* MEMOTE for standardized genome-scale metabolic

model testing. *Nat Biotechnol* 2020;**38**:272–6.

34. Thiele I, Palsson BØ. A protocol for generating a high-quality genome-scale metabolic reconstruction. *Nat Protoc* 2010;**5**:93–121.

35. Fritzemeier CJ, Hartleb D, Szappanos B *et al.* Erroneous energy-generating cycles in published genome scale metabolic networks: Identification and removal. Maranas CD (ed.). *PLoS Comput Biol* 2017;**13**:e1005494.

36. Maarleveld TR, Khandelwal RA, Olivier BG *et al.* Basic concepts and principles of stoichiometric modeling of metabolic networks. *Biotechnology Journal* 2013;**8**:997–1008.

37. Vinay-Lara E, Hamilton JJ, Stahl B *et al.* Genome –Scale Reconstruction of Metabolic Networks of Lactobacillus casei ATCC 334 and 12A. Andersen MR (ed.). *PLoS ONE* 2014;**9**:e110785.

38. Rousseeuw PJ. Silhouettes: A graphical aid to the interpretation and validation of cluster analysis. *Journal of Computational and Applied Mathematics* 1987;**20**:53–65.

39. Hawkey J, Vezina B, Monk JM *et al.* A curated collection of *Klebsiella* metabolic models reveals variable substrate usage and gene essentiality. *Genome Res* 2022:genome;gr.276289.121v2.

40. Baroukh C, Cottret L, Pires E *et al.* Insights into the metabolic specificities of pathogenic strains from the *Ralstonia solanacearum* species complex. Moleleki L (ed.). *mSystems* 2023:e00083-23.

41. Zboralski A, Biessy A, Savoie M-C *et al.* Metabolic and Genomic Traits of Phytobeneficial Phenazine-Producing *Pseudomonas* spp. Are Linked to Rhizosphere Colonization in *Arabidopsis thaliana* and *Solanum tuberosum*. Vieille C (ed.). *Appl Environ Microbiol* 2020;**86**:e02443-19.

42. Sayers EW, Bolton EE, Brister JR *et al.* Database resources of the national center for biotechnology information. *Nucleic Acids Research* 2022;**50**:D20–6.

43. Zimmermann J, Kaleta C, Waschina S. gapseq: informed prediction of bacterial metabolic pathways and reconstruction of accurate metabolic models. *Genome Biol* 2021;**22**:81.

44. Vezina B, Watts SC, Hawkey J *et al. Bactabolize: A Tool for High-Throughput Generation of Bacterial Strain-Specific Metabolic Models*. elife, 2023.

45. Nickel S, Steinhardt C, Schlenker H *et al.* IBM ILOG CPLEX Optimization Studio—A primer. *Decision Optimization with IBM ILOG CPLEX Optimization Studio*. Berlin, Heidelberg: Springer Berlin Heidelberg, 2022, 9–21.

46. Hyatt D, Chen G-L, LoCascio PF *et al.* Prodigal: prokaryotic gene recognition and translation initiation site identification. *BMC Bioinformatics* 2010;**11**:119.

47. Cock PJA, Antao T, Chang JT *et al.* Biopython: freely available Python tools for computational molecular biology and bioinformatics. *Bioinformatics* 2009;**25**:1422–3.

48. Liao Y-C, Huang T-W, Chen F-C *et al.* An Experimentally Validated Genome-Scale Metabolic Reconstruction of *Klebsiella pneumoniae* MGH 78578, *i* YL1228. *J Bacteriol*

2011;**193**:1710–7.

49. Peyraud R, Cottret L, Marmiesse L *et al.* A Resource Allocation Trade-Off between Virulence and Proliferation Drives Metabolic Versatility in the Plant Pathogen Ralstonia solanacearum. Desveaux D (ed.). *PLoS Pathog* 2016;**12**:e1005939.

50. Nogales J, Mueller J, Gudmundsson S *et al.* High‐quality genome‐scale metabolic modelling of *Pseudomonas putida* highlights its broad metabolic capabilities. *Environ Microbiol* 2020;**22**:255–69.

51. Jain C, Rodriguez-R LM, Phillippy AM *et al.* High throughput ANI analysis of 90K prokaryotic genomes reveals clear species boundaries. *Nat Commun* 2018;**9**:5114.

52. Li F, Cheng CC, Zheng J *et al.* Limosilactobacillus balticus sp. nov., Limosilactobacillus agrestis sp. nov., Limosilactobacillus albertensis sp. nov., Limosilactobacillus rudii sp. nov. and Limosilactobacillus fastidiosus sp. nov., five novel Limosilactobacillus species isolated from the vertebrate gastrointestinal tract, and proposal of six subspecies of Limosilactobacillus reuteri adapted to the gastrointestinal tract of specific vertebrate hosts. *International Journal of Systematic and Evolutionary Microbiology* 2021;**71**, DOI: 10.1099/ijsem.0.004644.

53. Kristjansdottir T, Bosma EF, Branco dos Santos F *et al.* A metabolic reconstruction of Lactobacillus reuteri JCM 1112 and analysis of its potential as a cell factory. *Microb Cell Fact* 2019;**18**:186.

54. Frese SA, Benson AK, Tannock GW *et al.* The Evolution of Host Specialization in the Vertebrate Gut Symbiont Lactobacillus reuteri. Guttman DS (ed.). *PLoS Genet* 2011;**7**:e1001314.

55. Wegmann U, MacKenzie DA, Zheng J *et al.* The pan-genome of Lactobacillus reuteri strains originating from the pig gastrointestinal tract. *BMC Genomics* 2015;**16**:1023.

56. Lee J-Y, Han GG, Choi J *et al.* Pan-Genomic Approaches in Lactobacillus reuteri as a Porcine Probiotic: Investigation of Host Adaptation and Antipathogenic Activity. *Microb Ecol* 2017;**74**:709–21.

57. Li H. Minimap2: pairwise alignment for nucleotide sequences. Birol I (ed.). *Bioinformatics* 2018;**34**:3094–100.

58. Li H. New strategies to improve minimap2 alignment accuracy. Alkan C (ed.). *Bioinformatics* 2021;**37**:4572–4.

59. Li H, Handsaker B, Wysoker A *et al.* The Sequence Alignment/Map format and SAMtools. *Bioinformatics* 2009;**25**:2078–9.

60. Zhao X, Gänzle MG. Genetic and phenotypic analysis of carbohydrate metabolism and transport in Lactobacillus reuteri. *International Journal of Food Microbiology* 2018;**272**:12–21.

61. Dimonaco NJ, Aubrey W, Kenobi K *et al.* No one tool to rule them all: prokaryotic gene prediction tool annotations are highly dependent on the organism of study. Marschall T (ed.). *Bioinformatics* 2022;**38**:1198–207.

62. Biessy A, Novinscak A, Blom J *et al.* Diversity of phytobeneficial traits revealed by whole-genome analysis of worldwide-isolated phenazine-producing *Pseudomonas* spp. *Environmental Microbiology* 2019;**21**:437–55.

63. Muscariello L, De Siena B, Marasco R. Lactobacillus Cell Surface Proteins Involved in Interaction with Mucus and Extracellular Matrix Components. *Curr Microbiol* 2020;**77**:3831–41.

64. MacKenzie DA, Jeffers F, Parker ML *et al.* Strain-specific diversity of mucus-binding proteins in the adhesion and aggregation properties of Lactobacillus reuteri. *Microbiology* 2010;**156**:3368–78.

65. Venter SN, Palmer M, Steenkamp ET. Relevance of prokaryotic subspecies in the age of genomics. *New Microbes and New Infections* 2022;**48**:101024.
